# Deciphering the History of ERK Activity from Fixed-Cell Immunofluorescence Measurements

**DOI:** 10.1101/2024.02.16.580760

**Authors:** Abhineet Ram, Michael Pargett, Yongin Choi, Devan Murphy, Markhus Cabel, Nont Kosaisawe, Gerald Quon, John Albeck

**Affiliations:** Department of Molecular and Cellular Biology, University of California, Davis

## Abstract

The Ras/ERK pathway drives cell proliferation and other oncogenic behaviors, and quantifying its activity *in situ* is of high interest in cancer diagnosis and therapy. Pathway activation is often assayed by measuring phosphorylated ERK. However, this form of measurement overlooks dynamic aspects of signaling that can only be observed over time. In this study, we combine a live, single-cell ERK biosensor approach with multiplexed immunofluorescence staining of downstream target proteins to ask how well immunostaining captures the dynamic history of ERK activity. Combining linear regression, machine learning, and differential equation models, we develop an interpretive framework for immunostains, in which Fra-1 and pRb levels imply long term activation of ERK signaling, while Egr-1 and c-Myc indicate recent activation. We show that this framework can distinguish different classes of ERK dynamics within a heterogeneous population, providing a tool for annotating ERK dynamics within fixed tissues.

## Introduction

The RAS/ERK pathway directs multiple cellular behaviors and regulates tissue homeostasis (Lavoie et al., 2020). The terminal kinase in this pathway, Extracellular Signal-Regulated Kinase (ERK), is essential for cellular decisions to enter the cell cycle, migrate, or differentiate. Elevated ERK activity drives cancer and other diseases, and the quantitative strength and timing of ERK signaling play a critical role in disease progression and treatment. For example, individual cell fates can be altered by minor interruptions in ERK activity (Min et al., 2020). Additionally, residual ERK activity following targeted kinase inhibitor treatment determines therapeutic efficacy (Bollag et al., 2010), showing that proper measurement of pathway activation is an essential clinical parameter. Measuring phosphorylated ERK within patient tissue samples is a widely used diagnostic for cancer drivers and treatment potency. However, such methods to assay ERK activation are limited in their spatiotemporal resolution and quantitative accuracy.

The complexity of measuring ERK activity arises from the fact that the duration and amplitude of ERK activation influence the cellular interpretation of its signal. The pattern of activity influences expression of numerous target genes (ETGs), including the Immediate Early Genes (IEGs), both by activating mRNA production and by enhancing protein stability (Cook et al., 1999; Murphy et al., 2002, 2004; Nakakuki et al., 2010; Uhlitz et al., 2017). Advances in live-cell imaging and CRISPR tagging have allowed a higher-resolution view of how patterns of activation and deactivation (ERK dynamics) correlate with ETG expression at the single-cell level. Dynamic features of the ERK signal have been shown to differentially drive its target genes. ERK amplitude and duration are integrated over time by stabilization of Fra-1 protein levels, whereas c-Fos, Egr-1, and other genes are reported to respond maximally to intermediate frequencies of activation (Gillies et al., 2017; Saito et al., 2013; Wilson et al., 2017). In some systems, ERK pulsatility also correlates with proliferation and protection from apoptosis, while sustained activity correlates with cell cycle arrest (Aikin et al., 2020; Ender et al., 2022). These results highlight that the dynamic nature of ERK signaling can differentially activate genes, and therefore control cellular processes. Current assessments of Ras/ERK pathway activation measure levels of phosphorylated ERK (pERK) using antibody-based assays (Bollag et al., 2010; Escobar-Hoyos et al., 2020; Flaherty et al., 2010). However, a more informative measure of ERK activity would capture its dynamic history, enabling the observer to distinguish between cells with long-term constitutive activity or intermittent activation. Live-cell reporters provide a method to achieve this resolution in experimental settings, but cannot be used in humans and are often inaccessible in animal models. Furthermore, phospho-ERK concentration does not necessarily capture its activity within the cell, given the variable role of competing phosphatases (Gillies et al., 2020), and changes in ERK phosphorylation can occur rapidly within the cell, especially during pharmacological inhibition of the pathway or tissue isolation (Kleiman et al., 2011). Therefore, detection of pERK is an unreliable indicator of the longer-term activation history of ERK (Albeck et al., 2013). In this study, we explore the feasibility of estimating past ERK activity using antibody-based measurements of ETGs. Previous work has demonstrated that synthetic ETGs can capture ERK dynamics (Ravindran et al., 2022). While incorporation of such biosensors remains impractical, endogenous ETGs have a range of different sensitivities to dynamic ERK activity (Davies et al., 2020; Wilson et al., 2017), which could potentially be used to infer pathway activation history using fixed-cell measurements only. Such inferences could be used in biopsy tissues to infer the dynamic nature of ERK activity within tumor tissue. Oncogene induced ERK activation has been shown to be distinct from normal physiological patterns, and is often sustained (Aikin et al., 2020; Bugaj et al., 2018), thus knowledge of the types of signaling found in a tissue can be informative about the source of stimulating activity. Moreover, the duration of signaling suppression by inhibitors is of high interest, therefore this could be a way of assessing the efficacy of a treatment over a long window of time.

To date, most studies of ETGs have considered how changes in ERK signaling dynamics impact the expression of a given ETG. We pose the reverse question: can an individual cell’s ETG expression profile be decoded to infer the history of ERK activation? Furthermore, what are the best quantitative indicators of ERK activation, and how can the strength, duration, and frequency of ERK activation be predicted? To investigate these questions, we used a live-cell biosensor of ERK in combination with cyclic immunofluorescence for ETGs and other proteins regulated by ERK, including the canonical ETGs Egr-1, Fra-1, c-Jun, c-Myc, c-Fos, and phosphorylated proteins such as pERK, pc-Fos, and pRb (a downstream marker of ERK-dependent cell cycle entry; for convenience we collectively refer to all of these markers as ETGs). Using statistical models and machine learning to predict ERK activity features based on the expression of each protein, we find that each gene product reports ERK history with a different memory span. Of the measurements used, long-term, average ERK activity is predicted best by levels of Fra-1, while short term, recent ERK activation is predicted best by levels of Egr-1 and c-Myc. Lastly, we tested the limits of our method by mathematical simulations of ERK driven gene expression, finding that in theory, static immunofluorescence measurements can well recapitulate dynamic activation history with as few as 16 targets.

## Results

### A dataset linking live-cell ERK activity to ERK target immunofluorescence

To create a dataset that enables correlation of ERK activation to downstream target expression and modification, we first collected live ERK activity measurements in response to differential activation of the RAS/MAP Kinase pathway. We used EKAR 3.5, a calibrated, FRET-based biosensor of ERK activity, to measure single-cell activation in MCF10A mammary epithelial cells (Fig. 1a, S1a-d). With a series of Epidermal Growth Factor (EGF) concentrations, ERK activity was stimulated in a dose-dependent manner (Fig. 1b, S1e). To increase the diversity of activity patterns, we added MEK inhibitor (MEKi) at varying times after EGF stimulation, and included treatments where EGF was added at different timepoints of the experiment (Fig. 1c, S1e, Supplementary Table 1). The combination of MEKi treatments and the dose curve of EGF led to a broad spectrum of ERK signaling behaviors, with pulsatile activity varying in both duration and amplitude (Fig. 1d). Consistent with previous studies (Gillies et al., 2017; Ryu et al., 2015), we found that ERK activation is heterogenous from cell to cell within each dose of EGF stimulation.

**Figure 1:**
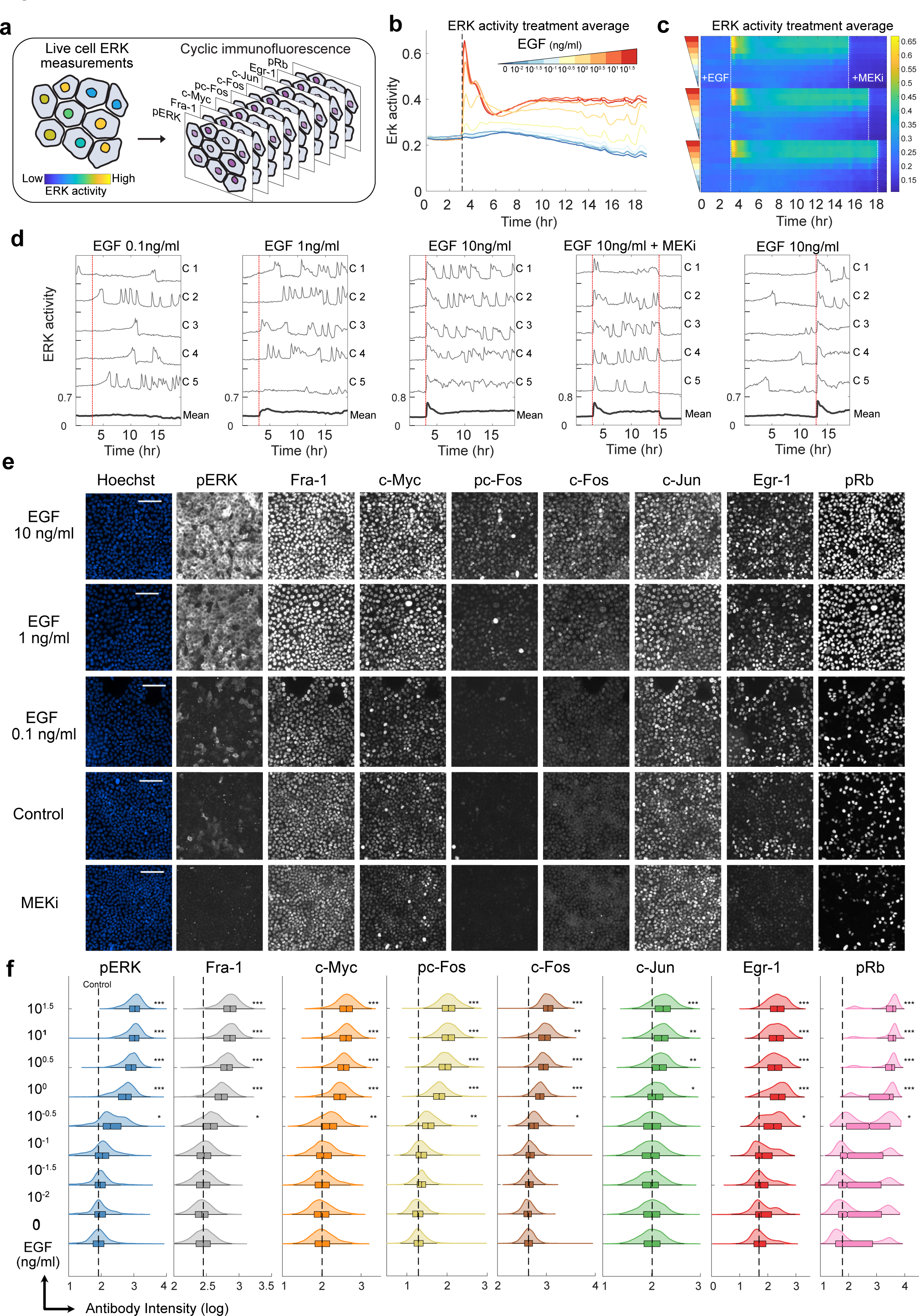
ERK activity and target genes are dose-responsive to Epidermal Growth Factor. **a** Schematic of the experimental method. Live cells were imaged in 96-well plates for 19 hours and immediately fixed. Plates were subsequently stained for antibody-based measurements. **b** Treatment average response measurements for live-cell ERK biosensor (EKAR) with increasing concentrations of EGF. Data are presented as the mean of each treatment (n_well replicates_ = 9-11 for each dose of EGF, 21 for control). **c** Treatment average response measurements depicted as a heatmap. Each row is the treatment average EKAR measurement (FRET measurements are indicated by color). EGF concentration indicated by colored triangles from Fig. 1b. MEKi = MEK inhibitor PD0325901 (100nM) (n_well replicates_ = 2-4 for each treatment). **d** Single-cell response plots to indicated treatment. Bold line indicates the average of all cells in one well of the treatment. **e** MCF10A cells immuno-stained with cyclic immunofluorescence. Each row depicts the same group of cells. Scale bar = 100 um. **f** Quantification of cyclic immunofluorescence measurements from listed EGF treatment. Dashed line indicates median of vehicle control condition (0 ng/ml EGF). Variance corrected t-tests were conducted by comparing each EGF treated condition to vehicle control n_replicates_ = 3. * p-val < 0.05, ** p-val < 0.005, *** p-val < 0.0005.

Immediately following live-cell data collection, we fixed the cells, and conducted cyclic immunofluorescence (4i) staining to measure levels of eight targets downstream of ERK (Fig. 1a, e, Supplemental Movie 1). This protocol was adapted from Gut et al. (Gut et al., 2018), and validated for our 96-well plate experiments (Fig. S2a-d). After quantifying antibody staining intensities, we found that most targets were dose responsive to EGF and suppressed by MEKi treatment (Fig. 1f, S1e). The one exception was c-Jun, which increased moderately with both MEK inhibition and EGF concentration, suggesting that its expression is not directly regulated by ERK activity in these cells.

We then analyzed the correlation between ERK activity and the expression of each target. To link live-cell ERK activity measurements with the respective 4i data for each cell, we aligned the corresponding image datasets and generated a heatmap arranged by the mean ERK activity measurement in each cell (Fig. 2a). While both ERK activity and 4i targets were variable across the data set, most of the 4i targets exhibited some discernible correlation with mean ERK activity, which was especially strong for Fra-1 and pRb. We calculated the Pearson correlation between ERK pulse features, such as frequency and duration for each cell, and each ETG measurement (Fig. 2b, c). The strongest correlations were between the sum of pulse duration to Fra-1 and pRb. Interestingly, Egr-1 was uniquely correlated with the average derivative of ERK activity, supporting the previous notion that Egr-1 selectively decodes pulsatile ERK activation (Saito et al., 2013). Of note, c-Jun had little to no correlation with any feature of ERK activation, implying again that its expression is not directly controlled by ERK in these cells and providing a useful negative control for subsequent analyses. We also performed a more granular time-sensitive analysis by calculating the Pearson correlation between each target and the EKAR FRET measurement at each timepoint of the live-cell movie (Fig. 2e). The correlation of Fra-1 and pRb was distributed across most of the time series (r= ∼0.5, 0.4, respectively) from the initial stimulus, apart from a period where ERK activity is weakest, about 2 to 4 hours after EGF addition. In contrast, c-Myc, c-Fos, and pc-Fos mildly correlate to ERK activity about 5 hours from prior to fixation (r= ∼0.3), and the correlation is highest (r= ∼0.55) during the last hour before fixation. As expected, pERK most correlates to ERK activity immediately prior to fixation (r= ∼0.6). Finally, Egr-1 correlates only to ERK activities 30 minutes to 1 hour prior to fixation (r: ∼0.5).

**Figure 2:**
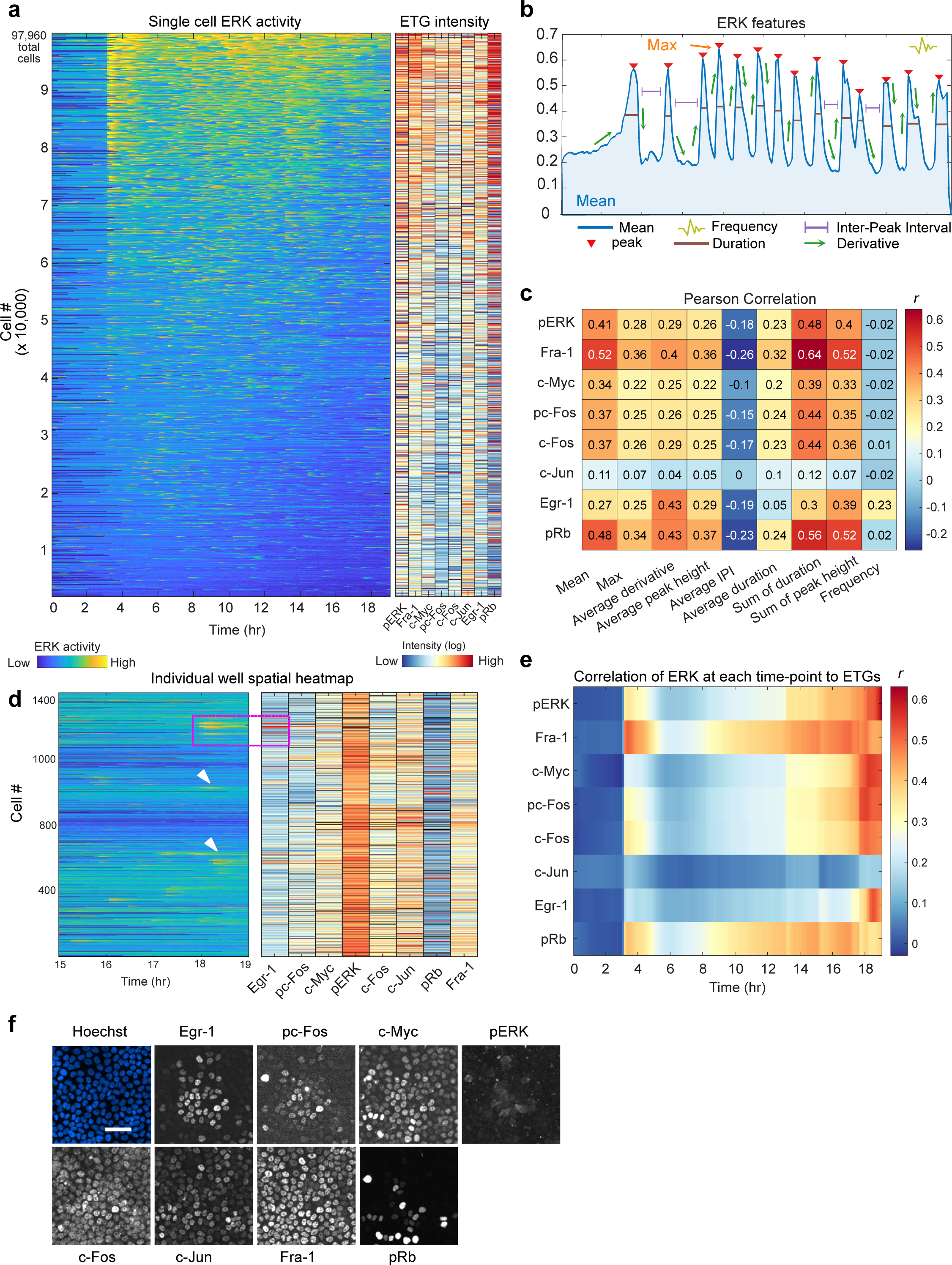
ERK target gene expression moderately correlates with features of ERK dynamics. **a** Single-cell heatmap for EKAR FRET measurements and corresponding ETG intensity, each row represents one cell (n_cells_ = 97,960, n_replicates_ = 3). ETG expression colored by log of antibody intensity from immunofluorescence measurements. **b** Features of ERK dynamics analyzed. Frequency was also calculated by estimating the mean normalized frequency of the power spectrum of the EKAR FRET measurement time series for each cell. **c** Pearson correlation (*r*) between each ERK feature and each cyclic immunofluorescence measurement, where single-cell values were used. **d** Spatial heatmap of EKAR (left) and ETG (right) measurements from a single well (control condition). Heatmap is organized by proximity of cells to each other so that neighboring cells in the well are plotted closer to each other in the heatmap. ETG colormap indicates relative log intensity of data within each column; outliers in pERK column skew colormap towards red. (black = NA). Magenta box indicates cells in Fig. 2f. White arrows indicate cells that recently activated ERK which resulted in higher Egr-1 expression (right). **e** Pearson correlation (*r*) between single-cell ETG measurements and the EKAR FRET measurement at each timepoint from the live-cell experiment. **f** Corresponding cells from magenta box in Fig. 2d. Scale bar = 50um.

To visualize spatial correlations of ERK-ETG signaling within the dataset, we plotted a spatial heatmap of signaling and gene expression, where cells within a single image are clustered in a heatmap visualization by proximity to one another (Fig. 2d). This analysis shows spatial ERK activation of groups of cells throughout the experiment. Interestingly, recent activation events are typically marked by strong Egr-1 expression within a local group of cells (Fig. 2f).

### Regression modeling of the ERK-ETG relationship predicts features of ERK dynamics

For a rigorous statistical analysis of the relationship between ERK activity and ETG expression, we performed cross-validated linear regression using the 4i measurements as predictors and ERK pulse features as response variables. We first created single predictor models to assess how well each target individually predicts each ERK feature in an individual cell. Analysis of the variance explained (R^2^) for each model confirms the results from the Pearson correlation analysis. Fra-1 and pRb best predict the sum of duration of ERK pulses (R^2^ of 0.42, 0.31, respectively), and the average ERK activity in each cell (R^2^ of 0.28 and 0.23) (Fig. 3a, b., S3a). These results suggest the duration of ERK activation seems to have a stronger influence on gene expression than the strength of the activation.

**Figure 3:**
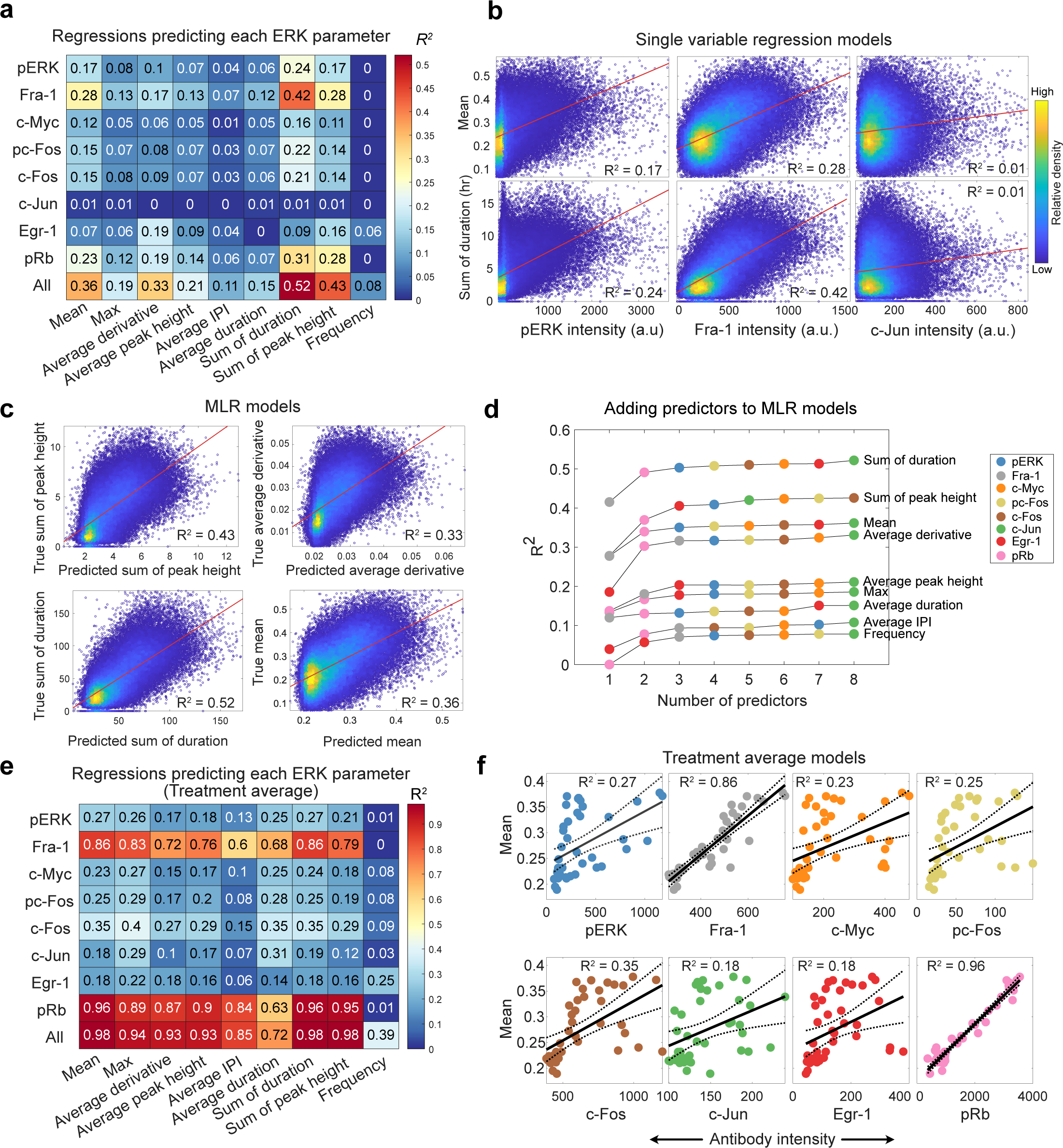
ERK target gene expression predicts history of ERK activation. **a** Single-cell regression showing the coefficient of determination (R^2^) of linear regression models which use ETGs to predict each ERK feature. 10-fold cross-validation was conducted to retrieve the best test-set model. This model was then fitted on the full dataset. **b** Scatter plots of single-cell regression models showing line of best fit. Color indicates relative density of the data. **c** Scatter plot showing each cell’s predicted (x-axis) vs true (y-axis) value in the multiple linear regression (MLR) models. **d** Results of adding predictors to MLR models. Color of each point indicates which predictor was added at each step. **e** Average values were calculated for all cells with the same treatment. These values were then used to fit regression models that predict each ERK feature using ETGs. **f** Scatter plots showing line of best fit and confidence intervals for treatment average regression models.

To assess whether the prediction models can be improved by considering multiple stains simultaneously, we generated several multiple linear regression (MLR) models using all 4i measurements as predictors at once (Fig. 3a, c). The ERK parameter with the highest variance explained was the sum of duration (R^2^ of 0.52) The models for derivative benefited the most from the multivariable models, however, they still only explained ∼33% of the variation in the data. c-Jun again served as a negative control, as those models did not explain any of the variation in ERK activity.

We then investigated which antibody combinations are most important in the MLR models. For each ERK feature, we created models that successively added predictors, and measured the resulting R^2^, and test-set error for each new predictor (Fig. 3d, S3b). For most ERK features, we found that the maximum R^2^ values can be achieved with just 2 to 3 predictors, where adding Fra-1 and pRb typically caused the highest improvement in R^2^ values and decrease in test-set error. The best model for the average derivative of ERK had a similar R^2^ with the model for the ERK mean (R^2^: 0.36, 0.33, respectively). The main distinction for the average derivative model is the strong contribution from Egr-1. To understand why adding more predictors does not improve models, we calculated the pairwise correlation between each 4i target, and found that many targets were moderately correlated with each other (Fig. S3c). The slight co-linearity between the predictors suggests that they share mutual information and thus explains why only a few predictors are needed to achieve the best possible models. These results indicate that ERK strength and duration are best inferred using Fra-1 and pRb, while ERK variability (derivative) are best inferred using Egr-1 and pRb. The fact that pRb predicts both long term ERK activation and variability suggests that Rb phosphorylation (cell cycle entry) is sensitive to different types of ERK activity (Min et al., 2020; Zwang et al., 2011).

We repeated the regression analysis on treatment averages to explore the difference between population and single-cell models. To calculate the treatment average for each measurement, we simply grouped cells with similar treatment conditions, and averaged their respective 4i measurements and ERK features. Overall, bulk models were superior to the single-cell models, as Fra-1 and pRb individually were excellent predictors for ERK dynamics (R^2^ > 0.7, Fig. 3e, f). Treatment average MLR models notably improved the predictions for average inter-peak interval, average duration, and frequency (R^2^ = 0.85, 0.72, 0.39, respectively). Fra-1 and pRb retained the most consistent relationship to ERK dynamics and were the most important predictors in all models. These results further solidify the importance of Fra-1 and pRb as markers for ERK activity. Furthermore, population average models reconcile the modest predictive power of the single-cell models and confirm the classical view that ERK determines gene expression.

Both the regression analysis and Pearson correlation indicate the pERK was not strongly correlated to long-term ERK activation in single cells. The poor relationship is partly due to the treatments that inhibit MEK at varying times after EGF addition, which lead to the virtually no pERK signal. When we remove these treatments from the analysis, the regression models notably improve for pERK, and slightly improve for other 4i measurements (Fig. S3d). These results indicate that pharmacological inhibition renders pERK an unreliable predictor of ERK histories, and that relying solely on pERK staining can lead to misinterpretations of pathway activation. Many studies assess the effect of pathway inhibitors using pERK staining; consequently, we argue other markers should be used. The Fra-1 and pRb models are robust to MEKi treatments, and therefore are the best predictors of long-term ERK activation.

### Neural network-based models of the ERK-ETG relationship reveals non-linear time dependence of ERK dynamics

While the previous models of featurized ERK activity provide interpretable correlations that help to understand the underlying biological process, they assume linearity and may not capture more complex relationships in the data. Additionally, some ERK parameters are correlated with each other, and other features of the time series may be missed. To examine the importance of the timing of ERK activation and identify which timepoints have the greatest impact on final expression levels, we trained a convolutional neural network (CNN) to use the ERK activity time series to predict expression levels of each ERK target in individual cells (Fig. 4a, top). As a comparison to the CNN, we also fit linear regression models (TS linear) using the values at each timepoint of the ERK time series as individual variables to predict final ERK target levels (Fig. 4a bottom left). Finally, we compared the performance of these time series-based models with that of ERK dynamics feature-based models (Featurized linear). These feature-based models used all nine featurized ERK measurements (i.e. mean/duration/frequency) to predict the expression of each ERK target (Fig. 4a bottom right).

**Figure 4:**
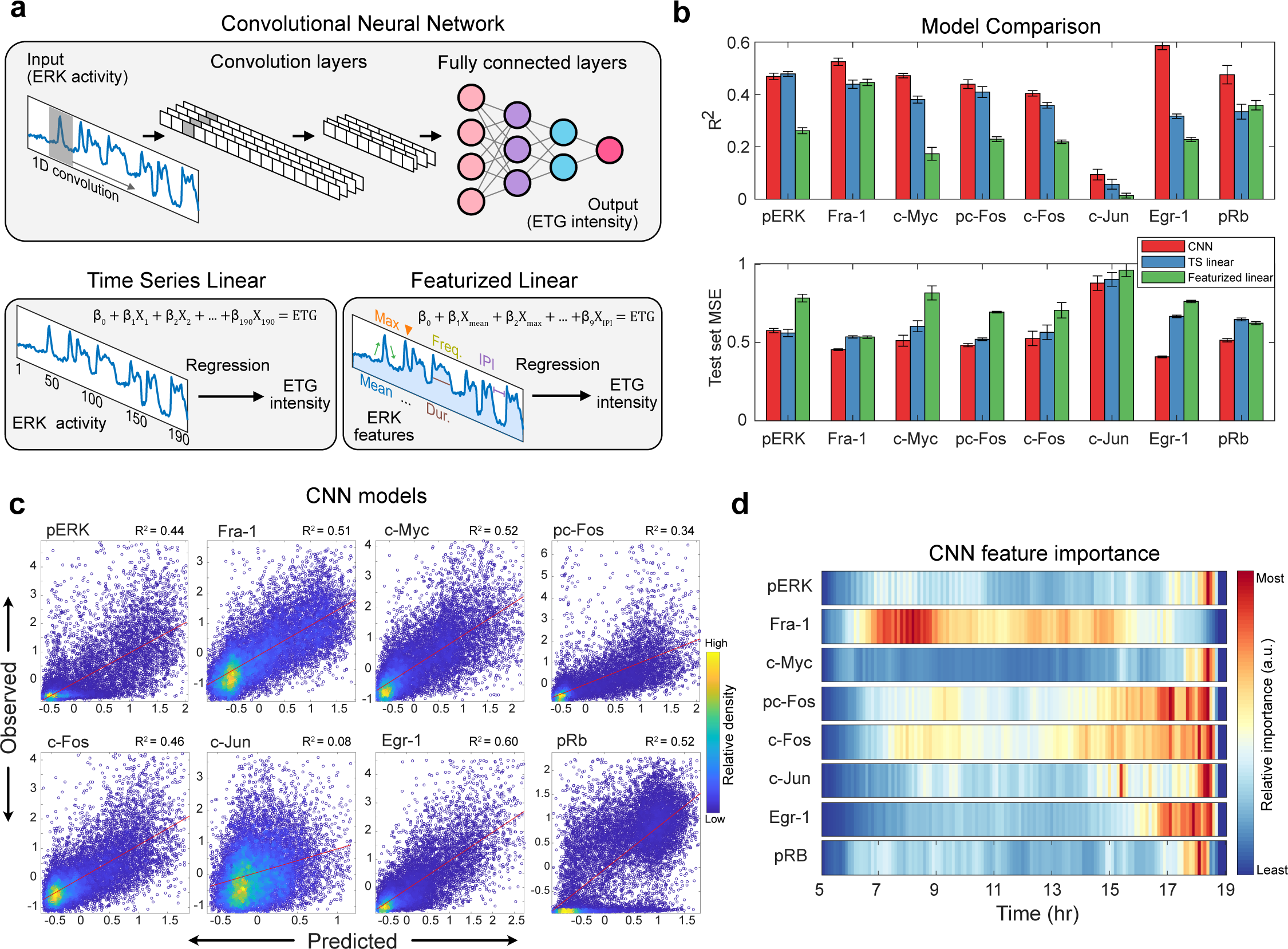
Convolutional neural network identifies non-linear signal transmission. **a** For each ETG, three types of prediction models were separately trained. Top: Simplified schematic of convolutional neural network architecture containing two convolutional layers and three fully connected layers. Bottom left: Multiple variable regression where ERK activity at each time point is considered as a predictor variable (TS linear). Bottom right: Multiple variable linear regression where nine features of ERK activity are considered as predictor variables (Featurized linear). **b** Top: Bar plot indicating R2 for three models used to predict ETG levels. Bottom: Bar plot indicating mean square error for three models used to predict ETG levels. Error bars represent standard error calculated using values from each fold of the 5-fold cross-validation partitions. **c** Scatter plot of the predicted and observed values of the CNN trained on all 190 timepoints (19 hr). The data represent standardized (z-scored) values. **d** Feature attribution heatmap showing the importance of each timepoint in the CNN model trained on 150 timepoints (15 hr). Color map represents relative values within each row.

We found that the CNN achieved the highest performance in predicting all ERK targets, except for pERK (Fig. 4b). To account for overfitting, we calculated the mean squared error (MSE) on unseen data (test set) and the CNN exhibited the least error for all targets, except for pERK (Fig. 4b bottom). Although the CNN yielded better performance for most targets, a significant amount of variance is still not captured by the model (Fig. 4c). Notably, the CNN models for Egr-1 and pRb explained much more variance than linear regression models of other targets, implying that Egr-1 and pRb likely respond to ERK activation with significant non-linearity. Finally, for many 4i targets, the featurized linear models underperformed the other two methods, both in R^2^ and test set error, indicating that the featurization method often does not capture important aspects of ERK signaling that influence gene expression.

We then used the CNN model parameter weights (feature importance) to investigate which timepoints most influenced the final expression of each target. However, feature importance across much of the time series was overshadowed by a strong correlation to the initial stimulus response, which likely reflects a correlation with the treatment delivered rather than direct biochemical regulation of ETGs (Fig. S4a). Therefore, we limited the model to using only time points more than 5 hours after the initial treatment when training our time series models, which resulted in a minimal decrease of the CNN performance (Fig. S4b). A CNN trained on fewer time points further confirms our regression results, and captures nuances of the ERK-ETG relationship (Fig. 4d). The findings reinforce the observations from the linear regression analysis, and highlight the importance of considering both the timing and intensity of ERK activation in understanding how gene expression is regulated. Fra-1 is influenced by a wide time span with peak influence starting as early as 12 hours prior; the most recent two hours have little effect. c-Fos and pc-Fos are also influenced by time spans of more than six hours, but focused on the last two to four hours. Egr-1 is strongly influenced by ERK activation within the last two hours, while pERK, c-Myc and pRb are influenced strongly by the last hour of ERK activation. This alternate modeling approach confirms that each ETG is differentially sensitive to timing of ERK activity, and that in some cases, this relationship is not well characterized as a linear relationship.

### Classification models uncover prototypical patterns of ERK signaling with distinct gene expression profiles

Thus far, we have trained models that predict several continuous variables that represent ERK history; however, the application of these models is limited by the challenge of concurrently visualizing the predictions. Therefore, we demonstrate here how spatiotemporal ERK predictions can be represented in a concise and intuitive manner. To do so, we first used k-means clustering to group cells into similar response classes, or prototypes, of ERK activity. We clustered cells into five classes: low activation (cluster 1), recent deactivation (cluster 2), long term activation (cluster 3), mid-term activation (cluster 4), and recent activation (cluster 5) (Fig. 5a, S5a). Analysis of the 4i target expression levels in each cluster was consistent with our previous statistical models (Fig. 5b). Long-term activation led to the highest expression of pERK, Fra-1, and pRb, while low activation displayed the lowest for all targets. Cells with recent activation highly expressed Egr-1 and c-Myc.

**Figure 5:**
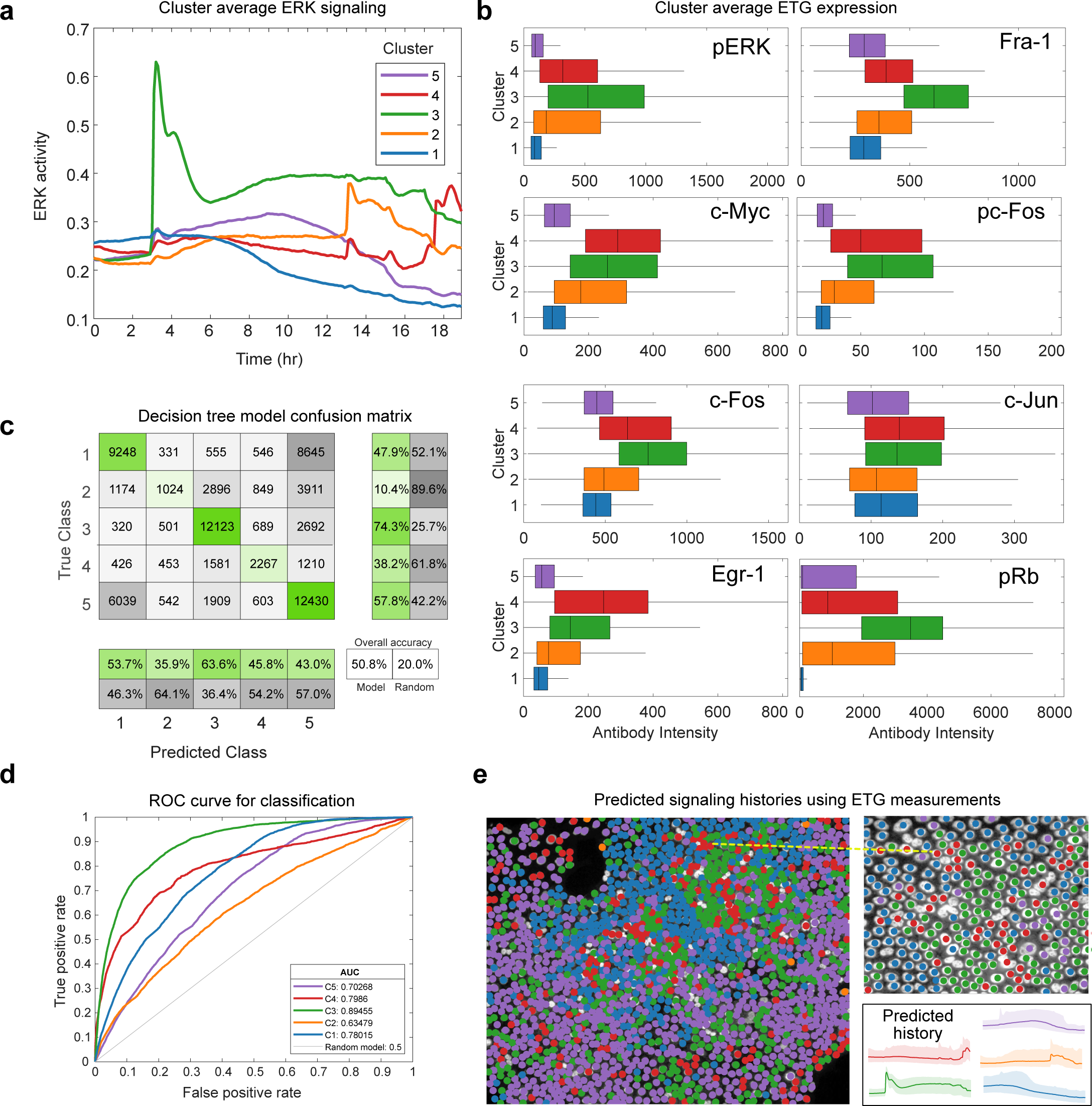
Annotating spatiotemporal ERK patterns in images using Decision Tree model. **a** Average ERK activity in each cluster identified by k-means clustering of EKAR time series data. Cells per cluster: C1_19325_, C2_9989_, C3_16325_, C4_6244_, C5_21523_. **b** Box plot showing median, quartiles, and range of ETG intensity in each cluster. **c** Confusion matrix showing the amount of accurate (green) and mis-classified (gray) cells in each class. Decision tree leaf size was optimized by cross-validation and collecting the leaf size with the minimum test-set error (129). **d** Receiver operating characteristic curves for each class in the decision tree model. **e** MCF10A cell stained with Hoechst (gray) overlayed with predicted signaling histories. Dark lines indicate the mean ERK activity for each cluster (as in Fig. 5a), and shaded regions indicate 25th and 75th percentiles.

We next trained a decision tree classifier that predicts prototypes of ERK signaling history using ERK target expression levels (Fig. 5c, d). The overall prediction accuracy of our model was 51% (compared to 20% for random selection), while individual class predictions varied in accuracy. Long-term activation class predictions were the most accurate (64%), and mid-term activation classifications were the least accurate (36%). These /findings indicate that long term and recent activation result in distinct patterns of the expressed genes we measured, while mid-term activation produces the highest variability in gene expression. The residual confusion in the classifier reflects that some classes are not well separated in the dataset, and that individual cells vary quite widely in their ERK activity (Fig. 5d, S5a). For classification, the predictor importance ranked pc-Fos as the most important predictor, followed by Fra-1 and pRb (Fig. S5b). This result indicates that, while pc-Fos may not explain a high amount of variance in ERK history, it carries particularly useful information for distinguishing among the five classes identified here. Finally, to simulate a potential use case with fixed tissue samples, we then used our classifier to predict ERK activity classes, and therefore histories, in cells from a single well in our dataset (Fig. 5e). Our analysis effectively quantifies the distinctiveness in gene expression associated with different ERK signaling prototypes and illustrates the utility of ETG stains in predicting the spatiotemporal signaling history of individual cells.

### Dynamical systems modeling of ERK-driven gene expression

To investigate the theoretical limits of predicting ERK dynamics from ETG levels, we extended an ordinary differential equation (ODE) model representing the regulation of ETGs (Davies et al., 2020; Gillies et al., 2017) (Fig. 6a). For a given ERK activity time series, the model simulates the mRNA and protein levels of a hypothetical ERK-responsive gene (sim-ETG). We constructed 1,000 hypothetical sim-ETGs by randomly assigning each one with different parameters values for mRNA degradation rate, protein degradation rate, phosphorylated protein degradation rate, protein dephosphorylation rate, negative feedback half-max concentration, and fractional expression at baseline (Fig. 6c, Supplementary Table 2). These 1,000 gene parameter configurations survey the parameter space with the goal of identifying sim-ETGs that capture different aspects of ERK signaling. Using 10,000 randomly selected live-cell ERK activity measurements from our experimental data, we simulated responses of all 1,000 sim-ETGs for each cell (Fig. 6b, S6A). Using the end point sim-ETG protein values (representing a fixed-cell 4i measurement of the hypothetical protein), we applied single variable regression modeling to characterize each sim-ETG’s capacity to predict ERK dynamics features. For predicting average ERK activity throughout the experiment, we found that 49% of sim-ETGs exhibited a R^2^ above 0.5 and over 100 were excellent predictors (R^2^ > 0.8) (Fig. 6d). For predicting the maximum activation and average pulse height, only 12% of sim-ETGs exhibited a R^2^ above 0.5, with a maximum R^2^ around 0.6 (Fig. S6c). Models for predicting dynamic ERK features like the frequency or the average derivative were overall worse than integrative features like the mean or sum of duration, reflecting that sim-ETGs under this model are variations on an integrator of ERK activity (Fig. S6c).

**Figure 6:**
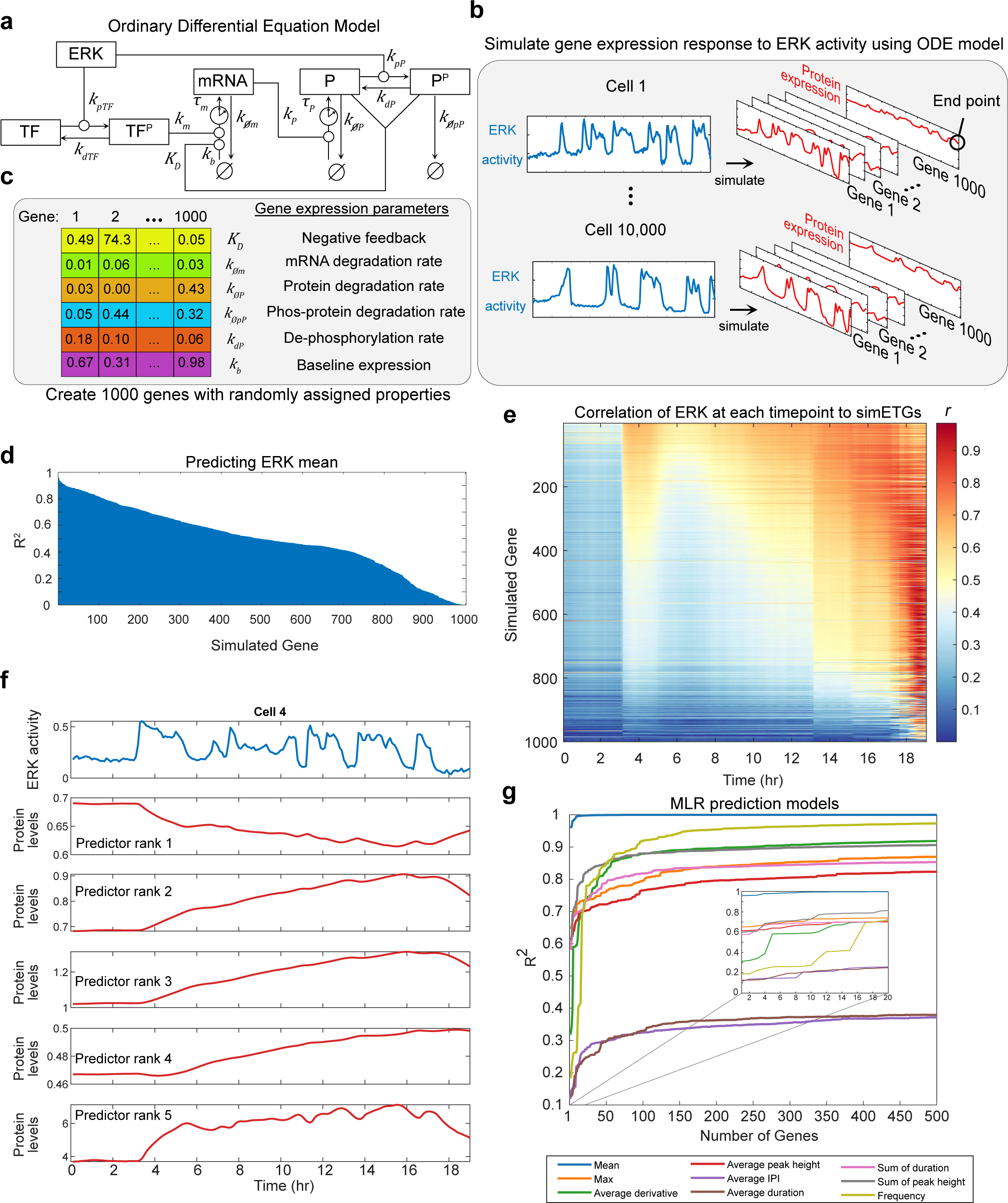
Mathematical model identifies limits of ERK activity prediction method. **a** Ordinary differential equation model representing ERK-dependent modification of a transcription factor (TF), expression of mRNA, and expression of a protein (P) product. Superscript P denotes phosphorylation of a molecule. Lowercase k’s indicate rate parameters, uppercase K indicates a dissociation constant for feedback effects. Clock icons indicate a time delay (τ). **b** 10,000 cells were randomly picked from our main dataset. We simulated the gene response of 1000 genes for each cell using our experimentally collected EKAR measurements. **c** 1000 simulated genes (sim-ETGs) were randomly assigned the listed rate parameters while other rate parameters in the model remained constant. **d** Coefficient of determination (R^2^) of single variable linear regression models using each sim-ETG to predict the average ERK activity in each cell. **e** Pearson correlation (*r*) between each sim-ETG (rows) measurement and the EKAR values at each timepoint from the live-cell experiment. **f** Top: one ERK activity trace from one cell in the dataset. Rest: gene expression response of the top 5 predictors of the mean ERK activity. **g** Multiple regression models fit to predict each ERK feature. For each ERK feature prediction model, a single sim-ETG was added as a predictor at each step. To determine the order of sim-ETGs to add, we performed single regression and ranked sim-ETG by their ability to individually predict each feature.

To visualize the gene expression response, we plotted a single cell’s ERK signal along with the response of the top five predictors of the mean (Fig. 6f). These response profiles show that both genes activated by or inhibited by ERK can serve as reliable predictors of ERK activity. While our experimental ETG measurements were selected based on known positive responders to ERK, 20% of sim-ETGs were negatively regulated by ERK (Fig. S6b); experimental prediction of ERK activity would likely be improved by including genes that are inhibited by ERK (Yamamoto et al., 2006). We then analyzed which gene parameters most influence how well an individual sim-ETG predicts mean ERK activity by examining the weights from a MLR model of sim-ETGs (Fig. S6d, e). Consistent with the known behavior of Fra-1, slow mRNA and phosphorylated protein degradation rates allow for accurate recording of the average ERK history.

Our 4i data analysis determined that while Fra-1 predicts long-term history, Egr-1 and c-Myc predict recent history. To examine this distinction in sim-ETGs, we calculated the correlation between the ERK activity at each timepoint and end-point protein expression (analogous to the experimental data in Fig. 2e). As expected, genes that predict mean ERK activity tend to be correlated with ERK activity over a wide time span, similarly to Fra-1. Those that are less effective at predicting mean are correlated with recent activation, behaving more like Egr-1 or c-Myc (Fig. 6e). Notably, no sim-ETG under this model was specifically predictive of intermediate timescales of activation (i.e. 5-10 hours prior to fixation).

Finally, to investigate how many gene measurements are required to accurately predict the different aspects of ERK signaling, we created MLR models which used many sim-ETGs at once to predict multiple ERK pulse features. These models greatly improved our predictions, as most explained 75 to 99% of the variance in the dataset (Fig. 6g). Of note, the derivative and frequency model predictions drastically improved as the number of predictors increased. This result was not obtained through overfitting, as the test set error of the models also decreased with more sim-ETGs (Fig. S6f). For most ERK features, between 16 to 20 sim-ETGs are required for obtaining good models (R^2^ ∼ 0.7) (Fig. 6g inset). From an experimental standpoint, these results demonstrate that predicting dynamic features of ERK is highly feasible, and depends largely on which gene products are measured. From a practical standpoint, measuring for 20 proteins using a multiplex staining protocol is readily achievable (Comandante-Lou et al., 2022; Stallaert et al., 2022). In all, the ODE model indicates that our ERK activation inference method is a feasible solution for fixed tissue analysis, and will benefit from further exploration of potential endogenous gene products to measure.

## Discussion

Here, we provide proof of principle that end-point ETG staining can be used to infer key aspects of long-term ERK activity within fixed cell samples. While differences in ETG activation by ERK were previously known, our analysis formalizes these differences and shows how quantitative models can be used to infer ERK’s activity history with single-cell resolution. The ETG measurements in these experiments provide information about two broad types of ERK behavior, long-term and short-term activation. Additionally, our model analysis of simulated ETGs demonstrates that additional measurements could even more finely resolve signaling patterns, such as intermittent pulses. The experimental and biological limits of these predictions remain to be established; however, this model framework can be used to estimate properties of ETGs that would optimally improve the measurement set.

While the main characteristics of ERK activity are captured by our models, a significant amount of unexplained variance in ERK activity in our analysis prompts the question of what other parameters can be used to improve ERK history predictions. While our dynamical systems modeling suggests that other direct ERK targets could be used, a recent study identified orthogonal markers of cellular state, such as Sec13 (a nuclear pore component) and Calreticulin (an Endoplasmic Reticulum-resident protein), that correlate highly with phosphorylated ERK (Kramer et al., 2022). These results imply that intrinsic cellular factors modify ERK signaling, and that these markers can improve our ERK prediction models. Furthermore, other cell state measurements that may improve predictions include protein translation rates, chromatin accessibility, or transcription factor availability.

The statistical models in this study were trained on data from diploid non-tumor mammary epithelial cells. Generalizing these methods for use in other cell lines or tissues will require similar datasets from a broad array of cellular settings because there are reported differences in some ETG responses among various cell types. For example, B-Raf inhibition disrupts signal transmission and alters the transcriptional response of c-Jun, Egr-1, and Cyclin-D1 (Bugaj et al., 2018). Additionally, different mutations in B-Raf can lead to induction or suppression of c-Jun (Comandante-Lou et al., 2022). Accordingly, prediction models should be trained on cell-line specific data, especially from cancer cells with different MAPK pathway mutations. Potentially, a much larger scope of experiment is needed to train a model to simultaneously capture many cell lines, for example by identifying either ETGs whose responses remain consistent, or additional targets that sufficiently reflect the cellular context. However, it is also possible that a small set of well-chosen measurements may be sufficient to generate a broadly useful model (Janes et al., 2005).

In the dataset presented here, phosphorylated-Rb was a surprisingly good predictor of both long term and short term signaling in both single-cell and population-level models. An important caveat about these results is that ERK biosensors (EKAR and ERK Kinase translocation reporter) have some sensitivity to cyclin-dependent kinases (CDKs). EKAR 3.5 in particular is sensitive to CDK1 during mitosis (Hirashima, 2022). Though mitotic events are typically much rarer than changes in ERK activity, some of the variation in our EKAR measurements likely arises from this CDK1 activity. Since EGF increases mitotic activity, pRb levels, and EKAR measurements, there will be some correlation between EKAR and pRb that is an artifact. This serves as a reminder that co-variance, or cross-talk, among measurements will bias these types of machine learning analyses, and should be carefully evaluated.

The results of this study suggest that the duration of signaling plays a stronger role in protein expression than the integrated activity. Such an effect could arise from saturation of a particular gene’s response to ERK, or from differences among genes in phosphatase specificity or other competing regulators. An alternative explanation for this result is that our ERK biosensor fails to capture the high ranges of ERK activation. However, we resolved this by calibrating the reporter to provide a linear readout of ERK substrate phosphorylation (Gillies et al., 2020). This leads to the possibility that ERK target genes have been selected for duration responders rather than signal integrators. Nonetheless, a fundamental question remains to be further explored: What is the biological resolution of strength, duration, and other features of ERK activity, with respect to gene expression? Our analysis provides some quantitative answers to how ERK activation patterns specify a subset of gene expression. Finally, the method can be employed further to more closely investigate the effect of ERK signaling on cell fate changes rather than gene expression.

Our method could be used to provide important details of ERK signaling within fixed tissue samples in a clinical setting. The ability to infer the long-term patterns of ERK activity in samples from patients treated with MEK, EGFR, or other targeted pathway inhibitors would provide a more reliable indication of the effectiveness of long-term ERK activity suppression, helping to reveal areas of drug resistance. By analogy, measurements of hemoglobin A1C provide a reliable indication of a patient’s time-averaged blood sugar that is useful in the clinical management of diabetes. We propose that inferring the longer-term characteristics of ERK activity will be of similar use in managing tumors that rely on aberrant signaling in this pathway. This strategy can further be applied to other dynamically regulated pathways implicated in disease such as metabolic, inflammatory, or stress response signaling. Both generalized and patient-specific models would allow for more accurate diagnoses and improve personalized medicine.

## Methods

### Reporter cell line generation

For these experiments, two stable cell lines were created by electroporating MCF10A 5e cells with the EKAR 3.5 construct on the piggyBAC transposase system (Pargett et al., 2017). Cells were selected with neomycin (250 μg/ml 2 weeks) until they were resistant to selection (∼2 weeks).

### Cell culture and media

All experiments were conducted with MCF10A 5e (Janes et al., 2010). Cells were maintained in DMEM/F12 supplemented with 5% horse serum, 20 ng/ml EGF, 10 μg/ml Insulin, 500 ng/ml hydrocortisone, and 100 ng/ml cholera toxin. 10cm plates were passaged approximately every four days and re-plated at a 1:10 dilution. Imaging experiments were conducted in custom DMEM/F12 lacking phenol red, riboflavin, and folate. This “imaging media” was supplemented with 500 ng/ml hydrocortisone, 17.5 mM glucose, 1 mM sodium pyruvate, 2 mM glutamine, 50μg/ml Penicillin/Streptomycin. Before plating cells for imaging experiments, 5 μlof Rat tail collagen was added to the middle of each well of a glass bottom 96-well plate (Cellvis) and incubated for 45 mins at 37°C. Cells were typsinized, plated at 6000 cells per well, and then incubated at 37°C for 45-60 mins. Growth media was then added, and the plate was left overnight in the incubator. The next day, immediately before the imaging experiment, the plate was washed 3x with imaging media, and the media was changed to imaging media. The experiment began one hour after this media change.

### Live cell microscopy and data acquisition

Prepared 96-well plates were imaged on a Nikon Ti-E inverted microscope with a stage-top incubator (37°C, 5% CO_2_). Coordinates within each well of the 96-well plate were imaged at 6 minute increments which were automated by the Nikon Elements AR software. Images were captured using an Andor Zyla 5.5 scMOS camera and a 20x/0.75 NA objective. Chroma #49001 (ET-CFP) and #49003 (ET-YFP) excitation/emission filter cubes were used for mTurquoise2 and YPet measurements, respectively. Further details are described in (Pargett et al., 2017). Coordinates of each acquisition area were saved for future imaging of immunostaining experiments.

### Cyclic immunofluorescence

Immediately after the final acquisition of the live cell experiment, cells were fixed in freshly prepared 12% paraformaldehyde for 10 min. Cells were then permeabilized with fresh, cold methanol for 10 mins (2 times total). Cells were then ready for iterative rounds of staining using a protocol adapted from (Gut et al., 2018). Briefly, the iterative protocol involves rounds of elution, blocking, primary staining, secondary staining, Hoechst staining, and finally image acquisition in a specific imaging buffer. Recipes for buffers are as follows: Elution buffer (0.5M Glycine, 3M Urea, 3M Guanidinium Chloride, 70mM TCEP), Blocking buffer (200mM NH_4_Cl, 300mM Maleimide, 2% BSA in PBS), primary/secondary staining buffer (200mM NH_4_Cl, 2% BSA in PBS), Hoechst-33342 stain (1:10,000 in PBS), and 4i imaging buffer (700 mM N-Acetyl Cysteine). Antibodies were incubated 24 to 48 hours from varying concentrations recommended by the manufacturer. The protocol was validated during the first replicate experiment to ensure that antibodies were properly eluted, data is shown in Fig. S2b-d. For the second and third replicate experiment, a visual inspection was completed prior to each round of staining to ensure proper antibody elution.

### Phos-tag western blotting

MCF10A 5e cells were plated on 6-well dishes the day before lysing. Cells were treated with EGF, PD0235901, or imaging media and lysed at the indicated timepoints. This procedure involved rinsing each well twice with ice cold PBS, cell scraping, and lysis with RIPA buffer (Sigma) with Halt protease inhibitor cocktail and 1 μM DTT. Cells were lysed at 80-90% confluency with a 50 μlof lysing buffer per well. 2 μlof each sample was then loaded in pre-cast phos-tag gels (Wako-Chem) and ran at 100V for 3 hours. The gel was chelated two times with transfer buffer and 10 mM EDTA for 15 minutes each and rinsed once more with just transfer buffer. Proteins were transferred overnight at 50V. The membrane was blocked with Li-COR Odyssey blocking buffer and blotted with anti-GFP antibody (24 hr incubation). The membrane was then blotted with Li-COR 800 anti-Mouse secondary antibody and imaged using a fluorescent scanner (Sapphire-Azure Biosystems). Intensities of the resulting phosphorylated EKAR 3.5 reporter and total EKAR 3.5 bands were measured in ImageJ.

### Image processing

Imaging data were saved as .nd2 files and accessed using the Bio-Formats toolbox for MATLAB (available from www.openmicroscopy.org/bio-formats), and processed with a custom MATLAB cell segmentation pipeline (Pargett et al., 2017). The procedure identified each cell’s nucleus using either EKAR 3.5 (live-cell) or Hoechst 33342 (IF) as a nuclear marker. The cytoplasm was defined as a ring around each cell’s nucleus. Background signal intensity was measured by imaging a well with no cells, but containing live-cell imaging media or 4i imaging buffer. Cell position tracking and linking was performed using uTrack 2.0 (Jaqaman et al,. 2008). The resulting single-cell data was filtered to remove cells with less than 15 hours of tracking data. FRET measurements of ERK activity for each cell were calculated with 1 – ((CFP/YFP) / R_p_), where CFP and YFP are the intensities of Cyan and YFP channels measured in each cell, respectively. R_p_ is the ratio of total power collected of CFP over YFP where the power of each channel is the integral of the spectral product of excitation intensity, filter transmittances, exposure time, fluorophore absorption and emission properties, and quantum efficiency of the camera (detailed in appendix of (Gillies et al., 2020)).

### Batch effect correction

To correct for batch effects in the immunofluorescence measurements across three replicates, we scaled measurements in logspace. For each 4i target, we calculated the median value for each treatment and matched identical treatments across replicates. These treatments included all EGF doses at timepoint 30, MEKi at timepoint 30, and imaging media control. We then took the log_10_ of these values and fit a linear model (equation 1):

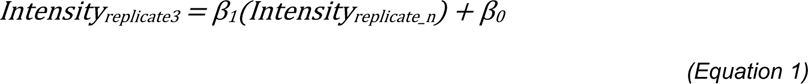

Where *Intensity_replicate_n_* represents log_10_ median values for either replicate 1 or replicate 2, *Intensity_replicate3_* represents the corresponding log_10_ median values for the third experimental replicate, and *β_0_* and *β_1_* are the scaling factors. These scaling factors were then used to correct all single-cell values for replicate 1 and 2. The corrected values were then returned to the linear scale by exponentiating.

### EKAR 3.5 Calibration

FRET measurements were calibrated to deliver a quantitative linear readout of ERK activity, as described previously (Gillies et al., 2020). Briefly, we used Phos-Tag immunoblotting to quantify the fraction of the EKAR 3.5 reporter that is phosphorylated in 3 concentrations of EGF (15 mins), phosphorylation inhibited (MEKi for 2 hours), and control conditions. These values were then linearly fit against the average EKAR 3.5 signal for the same conditions (equation 2).

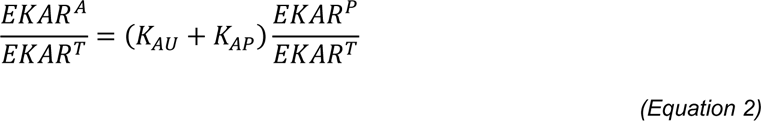

Where *EKAR^P^/EKAR^T^* represents the phos-tag ratio between phosphorylated and total reporter, and *EKAR^A^/EKAR^T^* represents the average FRET measurement in the corresponding condition.). *K_AU_* and *K_AP_* represent fractions of EKAR in the “associated” state when completely unphosphorylated and when completely phosphorylated, respectively. Single-cell FRET measurements *(f_A_)* were then used to estimate the concentration ratio of active ERK to the competing phosphatase(s) (equation 3). This ratio is the quantitative measure of ERK activity in a cell *(ERK^A^ / PPASE^A^)*.

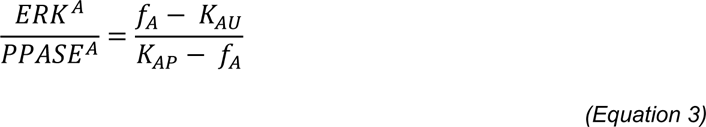

### Data analysis and regression modeling

Cells with less than 15 hours of data were removed prior to analysis, and cells out of the expected range of the FRET measurements were removed. FRET measurements were then adjusted using the reporter calibration model created from the phos-tag experiments. Thus, statistical models were created on cells that had complete EKAR and ETG measurements. Models were created using 10-fold cross validation. The data was randomly assigned to 10 groups, in which the 10^th^ group was held out of the model fitting procedure. The model was then tested against the 10^th^ group (test-set) to collect the test error (residual mean squared error, RMSE). This procedure is repeated for a total of 10 times to collect RMSE values from 10 test sets. The model that produced the lower test-set error was then refitted to the entire dataset to calculate the reported RMSE values.

### Pulse analysis and peak detection

The findpeaks function in MATLAB was used to find local maxima (peaks) for each cell’s ERK activity. Pulse features were then calculated based on the identified peaks. Frequency was calculated using the meanfreq function in MATLAB.This function estimates the mean normalized frequency of the power spectrum of each ERK activity trace.

### Statistical tests

For single-cell immunofluorescence data, each statistical comparison was made by t-test with unequal variances, and false discovery rate was controlled within each dataset via the Benjamini and Hochberg Step-Up procedure (α = 0.05). The variance for each experiment was determined from single-cell samples and added to variance across experiments. This corresponds to a linear error model: ε_i_ = ε_cell_ + ε_exp_, where the error (from the mean) of an individual cell ε_i_ equals the sum of the errors arising from cell-to-cell variation ε_cell_and from experiment variation ε_exp_.

### Spatial heatmap generation

Each cell’s time averaged coordinates were used to calculate the average Euclidean distance between each pair of cells within each well of the 96-well plate. Hierarchical clustering was performed on this distance matrix. The optimal leaf order was calculated by maximizing the sum of the similarity between adjacent leaves by flipping tree branches and without dividing the clusters. This order was then used to sort and display the live-cell and fixed-cell data.

### ETG prediction models and evaluation

For convolutional neural networks, we trained a CNN per ETG prediction. Each model consisted of 1) feature learning module and 2) prediction module. Feature learning module consists of 2 convolutional layers (16 channels and kernel size of 16) followed by an FC layer with size of 192 to match the initial input size. Prediction module consists of 2 FC layers (each size of 64 with relu activations) followed by a final linear FC layer that outputs a single ETG prediction. We trained one model per ETG for 100 epochs using Adam optimizer with learning rate of 0.001 and L2 regularization of 0.001. For the linear model, linear regression was implemented using sklearn python package with default parameters. The inputs were either the raw or featurized ERK activity for linear model or featurized linear model respectively. Evaluation on the model was performed using 5-fold cross validation with each fold roughly having the same representation from each well of origin and treatment.

#### Identification of significant input timepoints

For feature attribution approach, we used feature attribution, specifically Integrated Gradient (Sundararajan et al., 06--11 Aug 2017), to identify input timepoints that the model considers significant to prediction of ETG. Integrated Gradient was implemented using the python package Captum (Kokhlikyan et al., 2020). Feature attribution outputs score from each input time point to ETG per cell, which was averaged across cells for summarized visualization in the form of heatmap.

#### Backwards feature selection with timepoints after stimulation with CNN

To test the importance of the timepoints after the initial stimulation, we trained new CNN models to only use timepoints 2 hours after stimulation for ETG prediction. This model was trained on 15 hours of ERK activity data. The model and training setup used is identical to the setup used for the model with all the time points (19 hours of ERK activity data).

### Decision Tree Classifier

EKAR time series data were clustered into five groups using k-means clustering. Each group was assigned its class label. An optimized decision tree was fit using 8 ETG measurements to predict class labels of each cell. Leaf optimization was done by fitting multiple cross validated models and recording test-set error. This was done using MATLAB’s fitctree function. The model with the lowest test set error was chosen. Predictor Importance was estimated by summing the changes in the risk due to splits on every predictor and dividing the sum by the number of branch nodes– MATLAB predictorImportance function.

### Ordinary Differential Equation Modeling

The ODE model was adapted from Davies et al. 2020 (Davies et al., 2020). The model of ERK dependent gene expression (Equations 4, 5, 6, and 7) was constructed from a mass action approximation. This process is modeled in four steps (equation 4) phosphorylation of a transcription factor by ERK (TF^P^), (equation 5) transcription of target mRNA (mRNA), (equation 6) translation of target protein (P), and (equation 7) potential stabilization of target protein by ERK-dependent phosphorylation (P^P^). A regulatory term is included in the transcription process allowing negative feedback from target protein onto its own production. The model is formulated as a delay differential equation to account for the effective lag times of transcription and translation without explicitly addressing the complex processes involved.

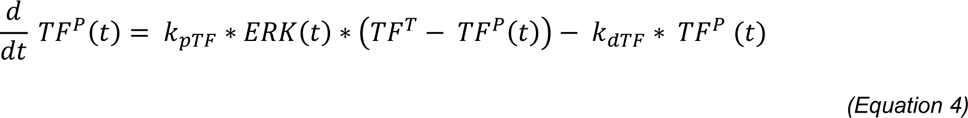

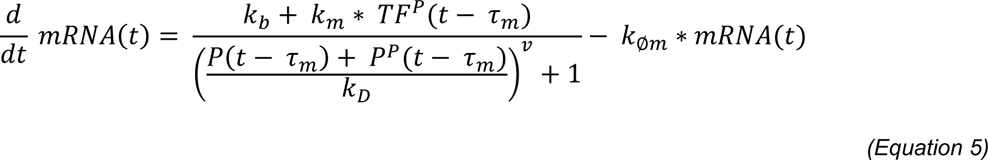

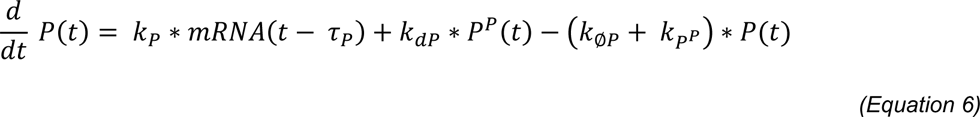

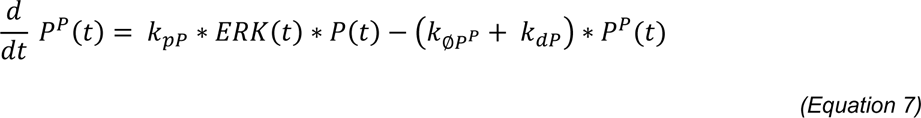

### Data availability statement

Raw and unprocessed data will be uploaded to BioStudies made publicly available. Processed data are uploaded as a .zip in the supplementary files of this manuscript.

### Code availability

Custom MATLAB code for data analysis used in this study are uploaded as a .zip file in the supplementary files of this manuscript.

### Author contributions

A.R., M.P., and J. A. conceptualized the study, interpreted data, and wrote the manuscript. A.R. conducted the imaging experiments and data analysis. M.P. created the ordinary differential equation model and the subsequent scripts that allowed for gene simulations. Y.C. trained the convolutional neural networks. D.M. assisted with western blot and image registration. M.C. assisted with cell culture. N.K assisted with data analysis.

## Acknowledgements

This work was supported by the National Institute of General Medical Sciences (R01GM115650 and R35GM139621 to JGA; T32GM007377 and 5R25GM056765 to AR) and the National Heart, Lung, and Blood Institute (R01HL151983 to JGA). We thank Carolyn Teragawa and Taryn Gillies for their helpful feedback on the manuscript.

---

**Supplementary Figure 1:**
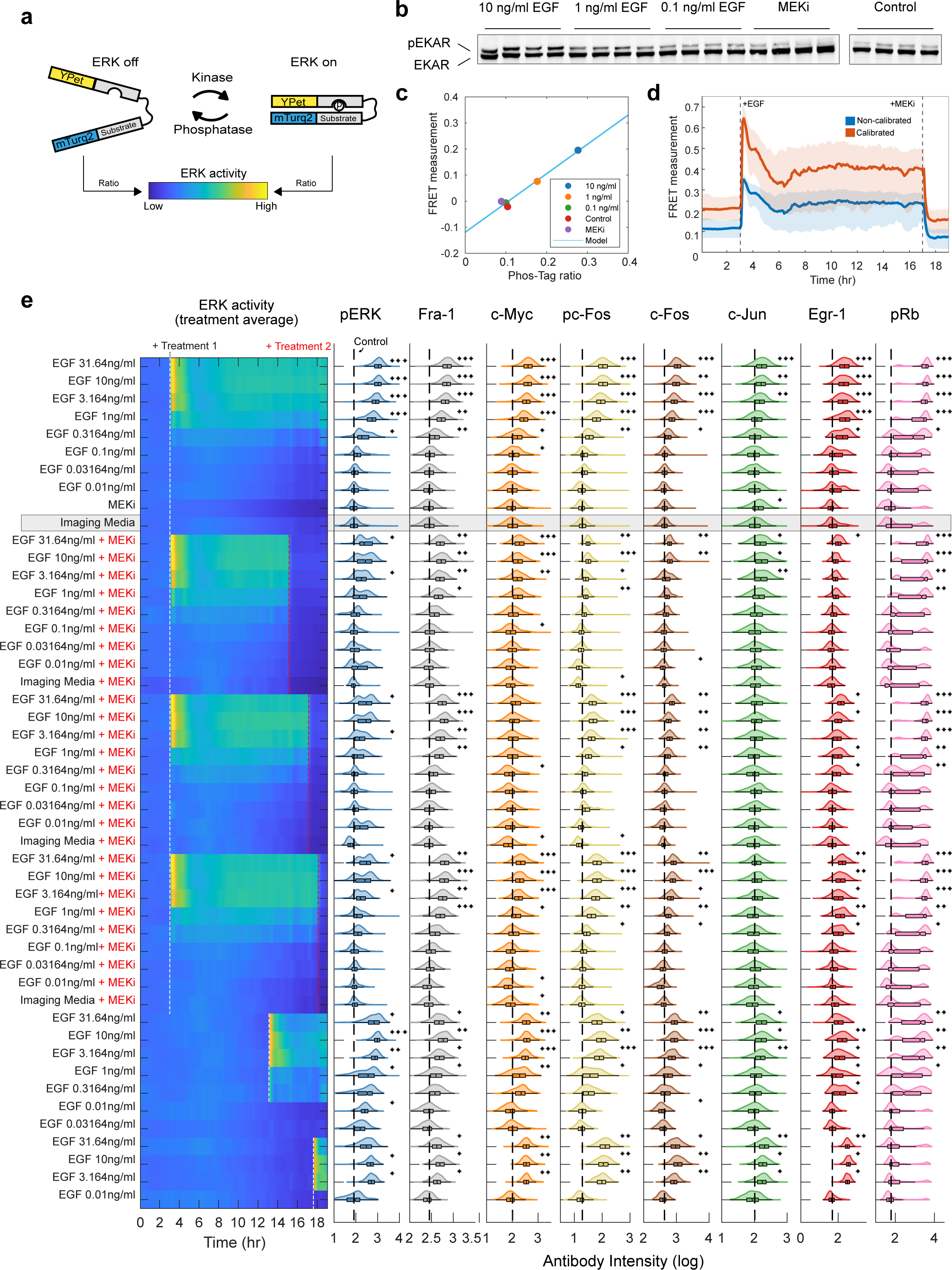
Live cell measurements with a calibrated ERK reporter followed by immunofluorescence. **a** Schematic of EKAR 3.5 FRET-based reporter. When ERK is inactive, mTurquoise2 and Ypet are distanced from each other. Active ERK binds the reporter substrate and induces a conformational change, bridging the two fluorescent proteins together. This causes a change in the ratio of mTurquoise2 and Ypet fluorescence intensities. **b** Phos-Tag immunoblot for phospho-EKAR 3.5 under 4 conditions that span the full range of ERK activity levels. Samples treated with EGF for 15 minutes, or MEKi for 2 hours. n_well replicates_ = 4 for each treatment. **c** Quantified ratio of the phosphorylated EKAR 3.5 over total EKAR 3.5 immunoblot intensities (x-axis). Y-axis represents the average live-cell FRET measurement in all cells within each treatment. FRET measurements were calculated at 15 minutes after EGF treatment, or 2 hours after MEKi. Each point represents the average of the 4 replicates. Model indicates the line of best fit. **d** Slope and intercept of the Phos-Tag model were used to calibrate the live-cell FRET measurements. **e** Left: Treatment average responses of EKAR biosensor data. Right: Histogram and box plot showing immunofluorescence quantifications for each treatment corresponding to the biosensor data. Box plot indicates median, quartiles, and range of the data. Dashed line indicates the median of the control (imaging media). Variance-corrected t-tests were conducted by comparing each EGF treated condition to vehicle control (imaging media) (n_replicates_ = 3). * p-val < 0.05, ** p-val < 0.005, *** p-val < 0.0005.

**Supplementary Figure 2:**
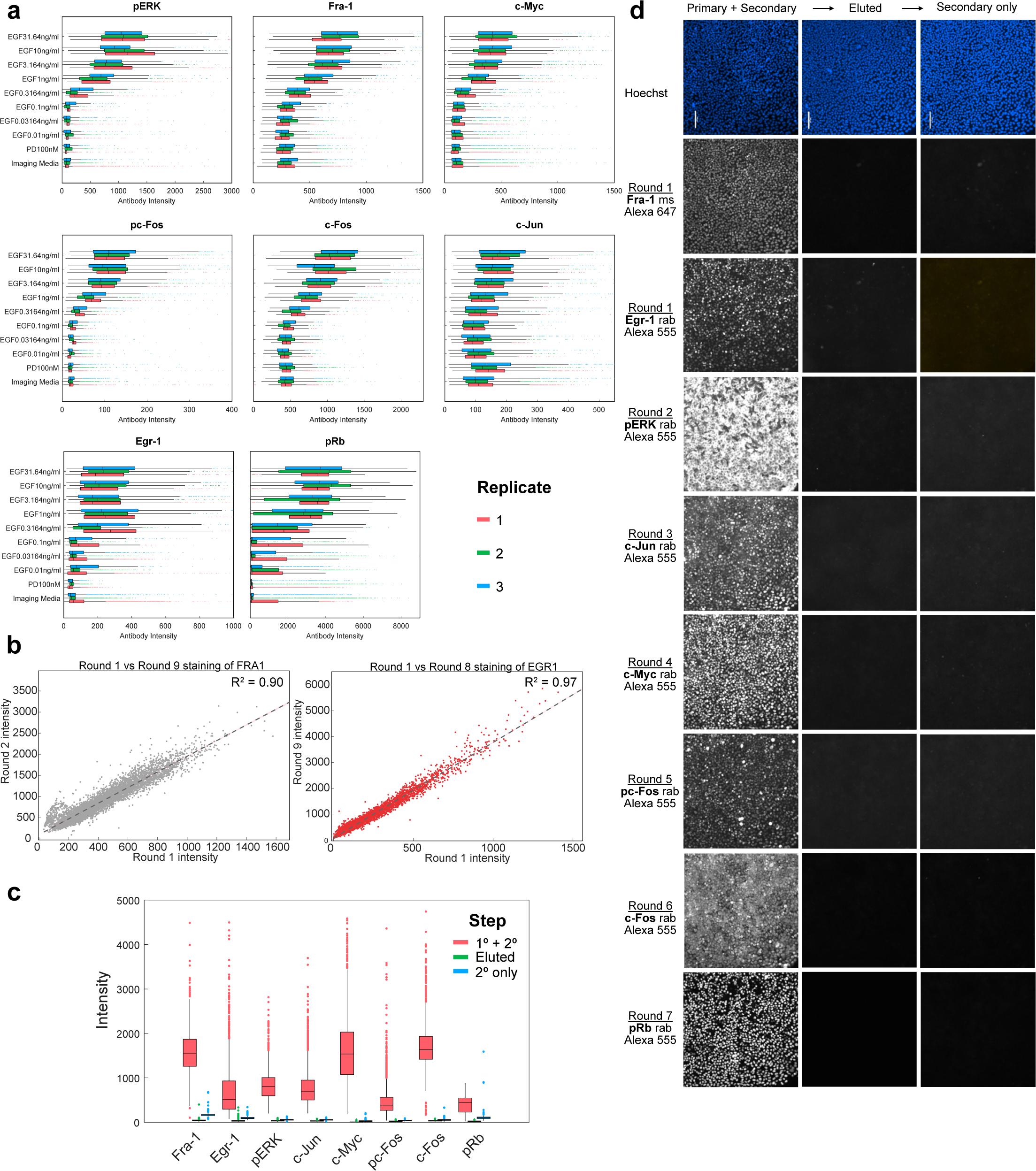
Batch effect correction and cyclic immunofluorescence protocol validation. **a** Box plot showing immunofluorescence quantifications for each treatment in each replicate experiment. Box plot indicates median, quartiles, and range of the data. Dots indicate outliers. n_replicates_ = 3. **b** Scatter plot of Fra-1 (left) and Egr-1 (right) intensity in the first round of staining vs. the ninth round from replicate plate 1. Data includes cells treated with EGF (all doses), imaging media, or MEKi. n_well replicates_ = 2 for each treatment. **c** Quantification of pixel intensities of cells in d. Box plot indicates median, quartiles, and range of the data. Dots indicate outliers. **d** Images of cyclic immunofluorescence rounds of staining. Cells were incubated with primary+secondary, eluted, and re-incubated with secondary only to ensure proper elution of the primary. Egr-1 and Fra-1 antibodies were both incubated together in round 1. Rab: anti-rabbit primary. Ms: anti-mouse primary. Cells treated with 31.64 ng/ml EGF. n_well replicates_ = 1. Five other wells treated with lower concentrations of EGF were also imaged and validated for proper elution (data not shown).

**Supplementary Figure 3:**
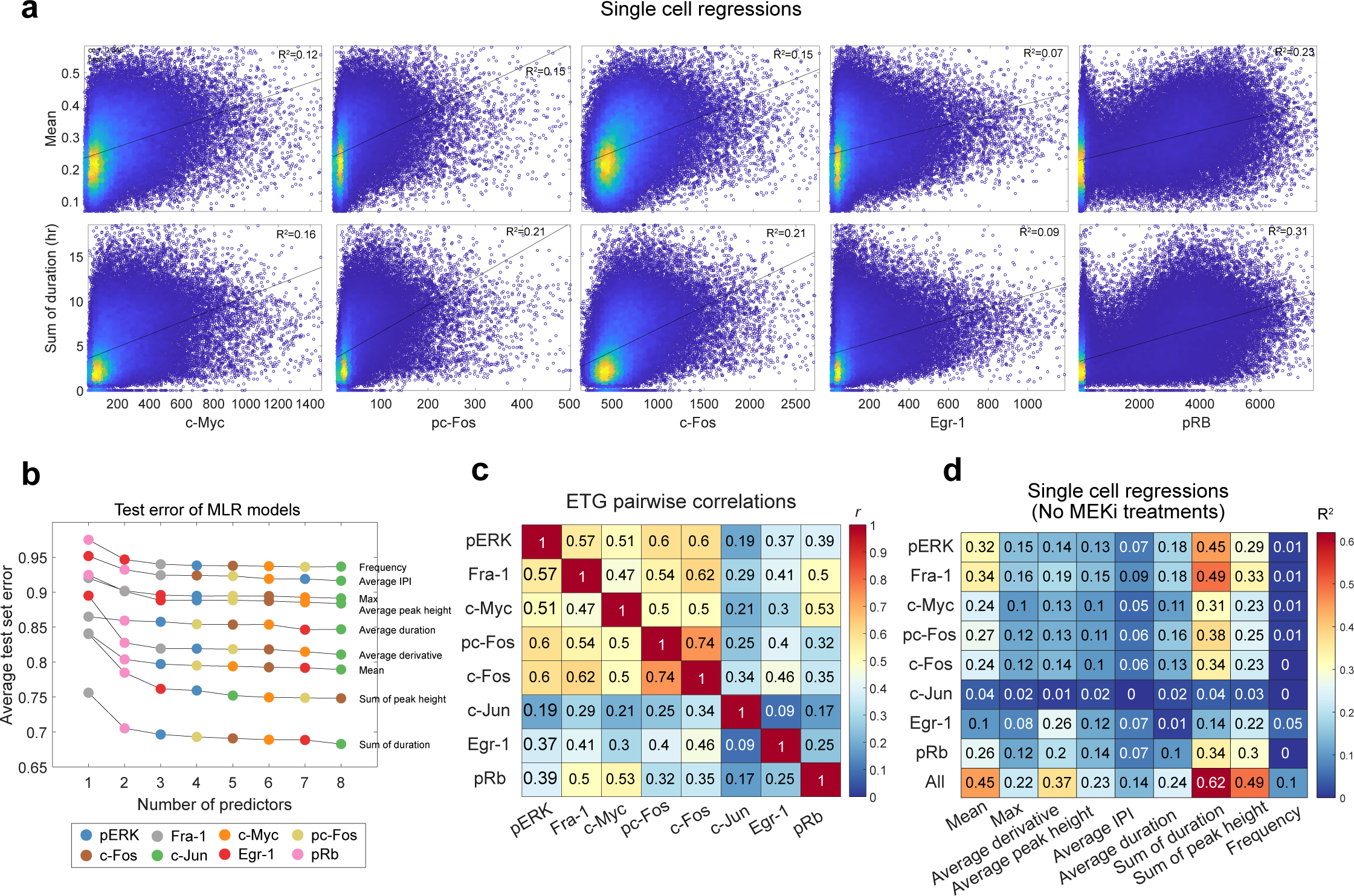
Regression modeling of ERK and ETGs. **a** Scatter plots and line of best fit for ETGs and ERK features. Color indicates relative density of data. **b** Test error (RMSE) of MLR models where additional predictors were added at each step. **c** Pearson correlation between each ETG. **d** Single variable regression models using single-cell data, cells treated with MEKi were removed from this analysis.

**Supplementary Figure 4:**
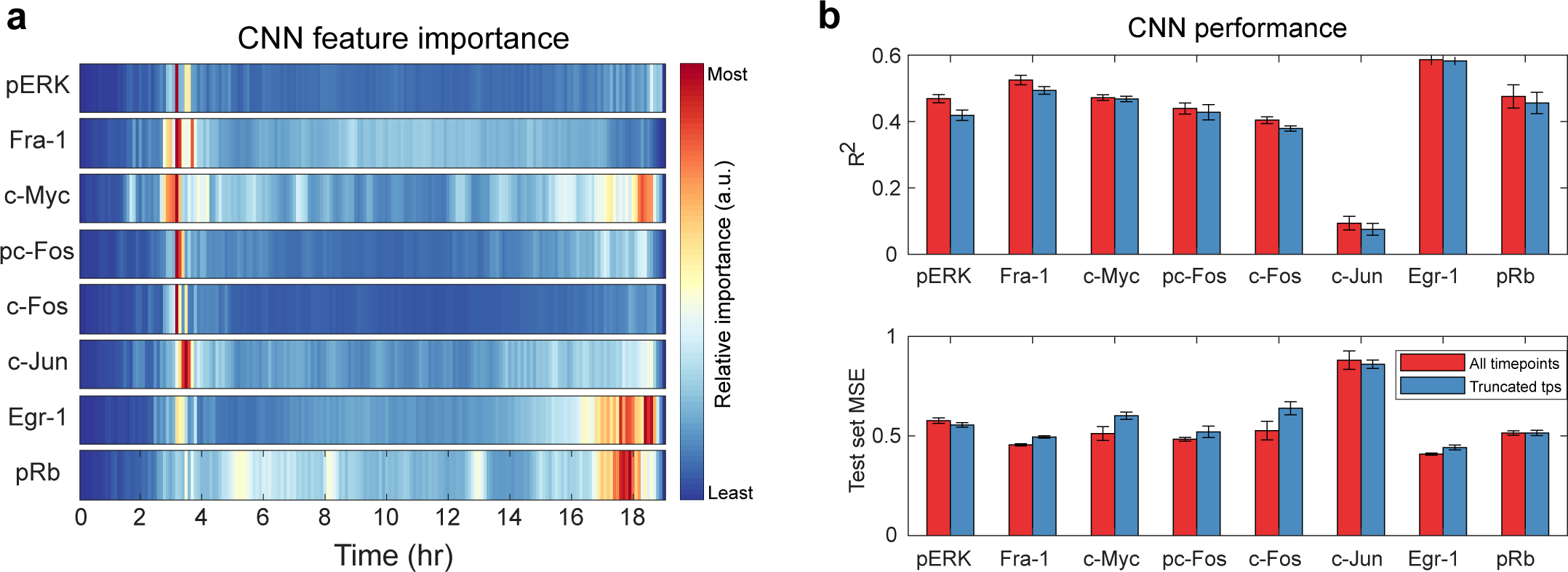
CNN feature importance is overshadowed by initial response. **a** Convolutional neural net feature importance of each timepoint in predicting levels of each ETG. Color map represents relative values within each row. **b** Comparing CNNs trained on 190 timepoints (19 hr) or 150 time points (15hr). Top: Bar plot of R2 value for predicting each ETG using k-fold cross-validation (k=5). For each ETG, data was partitioned into 5 groups. Within each k-fold, a training, test, and final set were created. Bar represents the average final set R2 value across all 5 groups. Error bars (Standard error) were calculated by dividing the standard deviation of R2 values for each ETG by the square root of five. Bottom: Test set mean squared error values for each ETG. Bar height and error bars were calculated as described above.

**Supplementary Figure 5:**
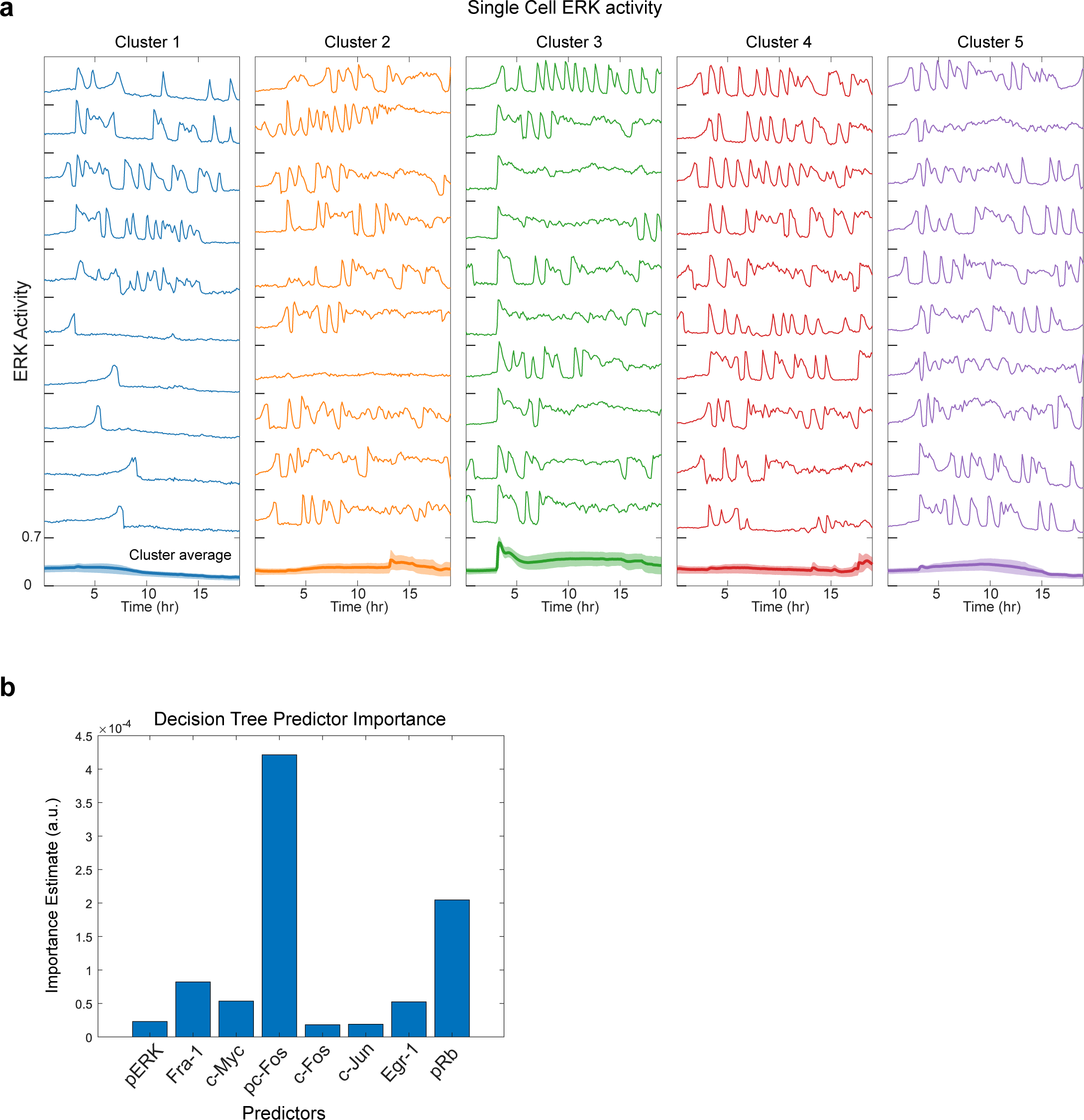
Single-cell variation with clusters. **a** ERK activity of 20 individual cells from each cluster identified by k-means clustering. Bottom line represents the cluster average, and the shading represents the 25th and 75th percentiles. **b** Predictor importance estimates for the decision tree classification model.

**Supplementary Figure 6:**
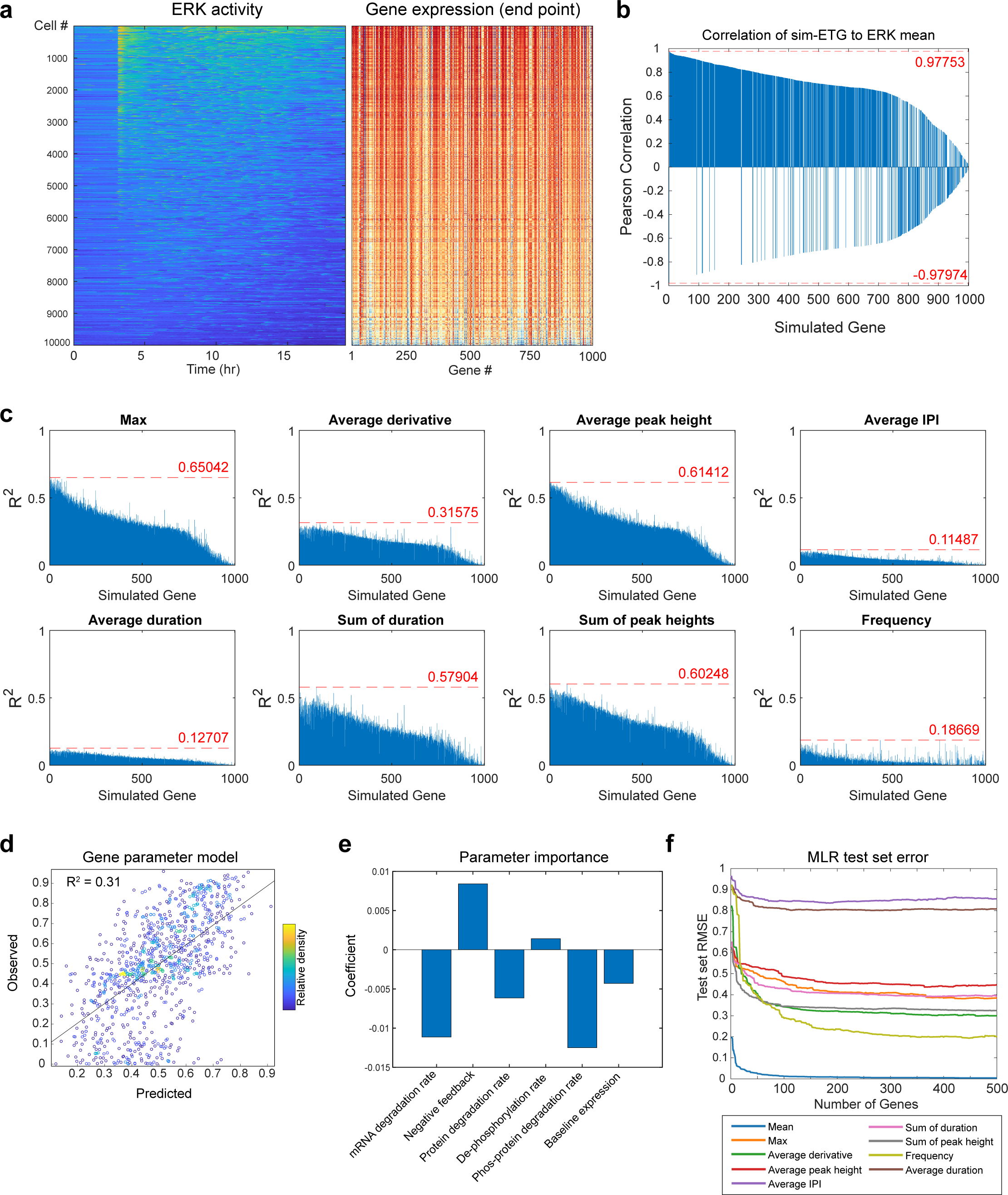
Ordinary differential equation modeling. **a** Left: single-cell ERK activity heatmap sorted by the mean of each cell (highest mean at the top). Right: Corresponding sim-ETG end point values. Color represents the relative expression within each column. n_cells_ = 10,000. n_genes_ = 1,000. **b** Pearson correlation between mean ERK activity and end-point gene expression. **c** R^2^ of single variable models using end point values of each sim-ETG to predict each ERK feature. Dashed line represents the maximum value. **d** Linear regression using gene parameter values to predict how well each gene tracks with average ERK activity. The model uses the negative feedback rate, mRNA degradation rate, protein degradation rate, phosphorylated protein degradation rate, de-phosphorylation rate, and fraction baseline to predict the R^2^ value from Fig. 6d. **e** Coefficient weights for linear regression in Fig. S6d. **f** Test set error (residual mean squared error) for each newly added gene in the prediction model. experiments.

**Supplementary table 1:** List of treatments/conditions along with the number of replicate wells and replicate experiments.

| Treatment1 | T1 Time (hr) | Treatment2 | T2 Time (hr) | Well replicates | Experimental replicates (days) |
| --- | --- | --- | --- | --- | --- |
| EGF 0.01ng/ml | 3 |  |  | 11 | 3 |
| EGF 0.03164ng/ml | 3 |  |  | 11 | 3 |
| EGF 0.1ng/ml | 3 |  |  | 9 | 3 |
| EGF 0.3164ng/ml | 3 |  |  | 10 | 3 |
| EGF 1ng/ml | 3 |  |  | 11 | 3 |
| EGF 3.164ng/ml | 3 |  |  | 11 | 3 |
| EGF 10ng/ml | 3 |  |  | 11 | 3 |
| EGF 31.64ng/ml | 3 |  |  | 11 | 3 |
| PD 100nM | 3 |  |  | 21 | 3 |
| Imaging media control | 3 |  |  | 32 | 3 |
| EGF 0.01ng/ml | 3 | PD 100 nM | 18 | 2 | 2 |
| EGF 0.03164ng/ml | 3 | PD 100 nM | 18 | 3 | 3 |
| EGF 0.1ng/ml | 3 | PD 100 nM | 18 | 3 | 3 |
| EGF 0.3164ng/ml | 3 | PD 100 nM | 18 | 3 | 3 |
| EGF 1ng/ml | 3 | PD 100 nM | 18 | 3 | 3 |
| EGF 3.164ng/ml | 3 | PD 100 nM | 18 | 3 | 3 |
| EGF 10ng/ml | 3 | PD 100 nM | 18 | 3 | 3 |
| EGF 31.64ng/ml | 3 | PD 100 nM | 18 | 3 | 3 |
| EGF 0.01ng/ml | 3 | PD 100 nM | 17 | 2 | 2 |
| EGF 0.03164ng/ml | 3 | PD 100 nM | 17 | 3 | 3 |
| EGF 0.1ng/ml | 3 | PD 100 nM | 17 | 3 | 3 |
| EGF 0.3164ng/ml | 3 | PD 100 nM | 17 | 3 | 3 |
| EGF 1ng/ml | 3 | PD 100 nM | 17 | 3 | 3 |
| EGF 3.164ng/ml | 3 | PD 100 nM | 17 | 3 | 3 |
| EGF 10ng/ml | 3 | PD 100 nM | 17 | 3 | 3 |
| EGF 31.64ng/ml | 3 | PD 100 nM | 17 | 3 | 3 |
| EGF 0.01ng/ml | 3 | PD 100 nM | 15 | 3 | 3 |
| EGF 0.03164ng/ml | 3 | PD 100 nM | 15 | 4 | 3 |
| EGF 0.1ng/ml | 3 | PD 100 nM | 15 | 4 | 3 |
| EGF 0.3164ng/ml | 3 | PD 100 nM | 15 | 4 | 3 |
| EGF 1ng/ml | 3 | PD 100 nM | 15 | 4 | 3 |
| EGF 3.164ng/ml | 3 | PD 100 nM | 15 | 4 | 3 |
| EGF 10ng/ml | 3 | PD 100 nM | 15 | 4 | 3 |
| EGF 31.64ng/ml | 3 | PD 100 nM | 15 | 4 | 3 |
| Imaging media control | 3 | PD 100 nM | 18 | 4 | 3 |
| Imaging media control | 3 | PD 100 nM | 17 | 2 | 1 |
| Imaging media control | 3 | PD 100 nM | 15 | 2 | 1 |
| EGF 31.64ng/ml | 17.5 |  |  | 2 | 2 |
| EGF 10ng/ml | 17.5 |  |  | 1 | 1 |
| EGF 0.01ng/ml | 17.5 |  |  | 1 | 1 |
| EGF 3.164ng/ml | 17.5 |  |  | 1 | 1 |
| EGF 0.01ng/ml | 13 |  |  | 1 | 1 |
| EGF 0.03164ng/ml | 13 |  |  | 1 | 1 |
| EGF 0.3164ng/ml | 13 |  |  | 2 | 2 |
| EGF 1ng/ml | 13 |  |  | 4 | 2 |
| EGF 3.164ng/ml | 13 |  |  | 2 | 2 |
| EGF 10ng/ml | 13 |  |  | 2 | 2 |
| EGF 31.64ng/ml | 13 |  |  | 2 | 2 |

**Supplementary table 2:** List of gene parameters in the ordinary differential equation model.

| Parameter | Description | Unit |
| --- | --- | --- |
| $TF^T$ | Total transcription factor concentration | nM |
| $k_{pTF}$ | ERK-dependent transcription factor phosphorylation rate | $\text{nM}^{-1}\text{min}^{-1}$ |
| $k_{dTF}$ | Transcription factor de-phosphorylation rate | $\text{min}^{-1}$ |
| $k_b$ | Baseline target mRNA transcription rate | nM/min |
| $k_m$ | ERK-dependent target mRNA transcription rate | $\text{min}^{-1}$ |
| $k_{\emptyset m}$ | Target mRNA degradation rate | $\text{min}^{-1}$ |
| $\tau_m$ | Transcription delay | min |
| $K_D$ | Negative feedback half-maximal concentration | nM |
| $\nu$ | Feedback Hill Coef. | - |
| $k_p$ | Target protein translation rate | $\text{min}^{-1}$ |
| $k_{\emptyset P}$ | Target protein degradation rate | $\text{min}^{-1}$ |
| $\tau_P$ | Translation delay | min |
| $k_{pP}$ | ERK-dependent target phosphorylation rate | $\text{nM}^{-1}\text{min}^{-1}$ |
| $k_{dP}$ | Target protein de-phosphorylation rate | $\text{min}^{-1}$ |
| $k_{\emptyset pP}$ | Phosphorylated target degradation rate | $\text{min}^{-1}$ |

**Supplementary table 3:** List of materials, software, and reagents used in the study.

| Reagent or Resource | Source | Identifier/RRID |
| --- | --- | --- |
| **Antibodies** |  |  |
| Anti-Fra-1, clone C-12 | Santa Cruz Biotechnology | sc28310; AB_627632 |
| Anti-Egr-1, clone 44D5 | Cell signaling | 4153; AB_2097035 |
| Anti-phospho-ERK p44/42 clone D13.14.4E | Cell signaling | 4370; AB_2315112 |
| Anti-cJun clone 60A8 | Cell signaling | 9165; AB_2130165 |
| Anti-c-Myc clone D84C12 | Cell signaling | 5605; AB_1903938 |
| Anti-phospho-c-Fos clone D82C12 | Cell signaling | 5348; AB_10557109 |
| Anti-c-Fos clone | abcam | ab190289; AB_2737414 |
| Anti-phospho-Rb (Ser807/811) clone D20B12 | Cell signaling | 8516; AB_11178658 |
| Goat anti-Rabbit IgG Alexa Fluor 647 | ThermoFisher | A-21245; AB_2535813 |
| Donkey anti-Rabbit IgG (H+L) Alexa Fluor 555 | ThermoFisher | A-31572; AB_162543 |
| IRDye 800CW Donkey anti-Mouse IgG | Licor | 926-32212; AB_621847 |
| **Chemicals, Peptides, and Recombinant Proteins** |  |  |
| Epidermal growth factor | Peprotech | AF-100-15 |
| PD0325901 | Selleck Biochemicals | S1036 |
| Cholera Toxin | Sigma-Aldrich | C8052 |
| Hydrocortisone | Sigma-Aldrich | H0888 |
| Insulin | Sigma-Aldrich | I9278 |
| Bovine Serum Albumin | Sigma-Aldrich | A7906 |
| Heat Inactivated Horse Serum | Life Technologies | 26050 |
| DMEM/F-12 1:1 | Life Technologies | 11320 |
| Neomycin |  |  |
| Collagen I, rat tail | Life Technologies | A10483-01 |
| L-Glutamine | Life Technologies | 25030-081 |
| Penicillin streptomycin | Life Technologies | 15070-063 |
| 0.25% Trypsin-EDTA | Life Technologies | 25200-056 |
| Tris Base | Fisher | BP152 |
| Glycine (Crystalline Powder) | Fisher | BP381 |
| Ponceau S solution, suitable for electrophoresis, 0.1% (w/v) in 5% acetic acid, 1L | Sigma-Aldrich | P7170-1L |
| Bromophenol blue | Sigma-Aldrich | B5525 |
| Dithiothreitol | Fisher | BP172 |
| Ammonium chloride | Sigma-Aldrich | 254134 |
| Maleimide | Sigma-Aldrich | 129585 |
| N-Acetyl-L-cysteine | Sigma-Aldrich | A7250 |
| TCEP hydrochloride | ApexBio | B6055 |
| Hoechst-33342 | Life Technologies | H3570 |
| Odyssey Blocking Buffer (PBS) | Licor | 927-40000 |
| Urea | Fisher | U15-500 |
| Guanidinium Hydrochloride | Fisher | BP178-500 |
| Phosphate-buffered saline, | Fisher | BP399-1 |
| Paraformaldehyde | ThermoFisher | 043368.9M |
| Glycine | Fisher | BP381-500 |
| Tween-20 | Fisher | BP337100 |
| Halt Protease inhibitor cocktail | ThermoFisher | 1861278 |
| **Experimental Models/Cell Lines** |  |  |
| Human: MCF-10A, clone 5E | Joan Brugge, Harvard Medical School | RRID:CVCL_0598 |
| **Recombinant DNA** |  |  |
| Plasmid: pPBJ-EKAR3.5nls-neo | Sparta et al. 2015 | Addgene # forthcoming |
| **Software and Algorithms** |  |  |
| NIS-Elements AR ver. 4.20 | Nikon | RRID:SCR_014329 |
| Bio-Formats ver. 5.1.1 (May 2015) | OME | RRID:SCR_000450 |
| uTrack 2.0 | (Jaqaman et al., 2008) | http://www.utsouthwestern.edu/labs/danuser/software/ |
| MATLAB 2020a | Mathworks | SCR_001622 |
| ImageJ (1.52p) | National Institutes of Health | RRID:SCR_003070 |
| Python (3.8.16) | Python Software Foundation | RRID:SCR_008394 |
| **Proofreading software** |  |  |
| Grammarly | Grammarly, Inc | www.grammarly.com/ |
| ChatGPT | OpenAI | www.openai.com |
| **Other** |  |  |
| Glass Bottom Plates, #1.5 cover glass | In Vitro Scientific | P24-1.5H-N, P96-1.5H-N |
| SuperSep Phos-tag gels (50 µmol/l), 12.5%, 17 wells | Wako-Chem | 195-17991 |

